# Accelerated human liver progenitor generation from pluripotent stem cells by inhibiting formation of unwanted lineages

**DOI:** 10.1101/174698

**Authors:** Lay Teng Ang, Antson Kiat Yee Tan, Matias Ilmari Autio, Joanne Su-Hua Goh, Siew Hua Choo, Kian Leong Lee, Jianmin Tan, Bangfen Pan, Jane Jia Hui Lee, Isabelle Kai Xin Yeo, Chloe Jin Yee Wong, Jen Jen Lum, Chet Hong Loh, Ying Yan Lim, Jueween Ling Li Oh, Cheryl Pei Lynn Chia, Angela Chen, Qing Feng Chen, Irving L. Weissman, Kyle M. Loh, Bing Lim

## Abstract

Despite decisive progress in differentiating pluripotent stem cells (PSCs) into diverse cell-types, the often-lengthy differentiation and functional immaturity of such cell-types remain pertinent issues. Here we address the first challenge of prolonged differentiation in the generation of hepatocyte-like cells from PSCs. We delineate a roadmap describing the extracellular signals controlling six sequential branching lineage choices leading from pluripotency to endoderm, foregut, and finally, liver progenitors. By blocking formation of unwanted cell-types at each lineage juncture and manipulating temporally-dynamic signals, we accelerated generation of 89.0±3.1% AFP^+^ human liver bud progenitors and 87.3±9.4% ALBUMIN^+^ hepatocyte-like cells by days 6 and 18 of PSC differentiation, respectively. 81.5±3.2% of hepatocyte-like cells expressed metabolic enzyme FAH (as assayed by a new knock-in reporter line) and improved short-term survival in the *Fah*^-/-^*Rag2*^-/-^*Il2rg*^-/-^ mouse model of liver failure. Collectively the timed signaling interventions indicated by this developmental roadmap enable accelerated production of human liver progenitors from PSCs.

## Introduction

A quintessential goal of regenerative medicine has been to manufacture various human cell-types in a dish from pluripotent stem cells (PSCs). Despite decisive progress, PSC differentiation into many lineages often takes weeks or months of *in vitro* differentiation; yields cell-types with limited functional maturity; and may produce heterogeneous populations containing other unwanted cell-types (Cohen and Melton, 2011). With regard to the first challenge (prolonged differentiation), one notion was that PSC differentiation might be an innately lengthy process. By contrast, PSC differentiation towards neuronal and mesodermal lineages can be considerably accelerated by precisely manipulating developmental signals in a temporally-dynamic fashion (Chambers et al., 2012; Loh et al., 2016; Qi et al., 2017). Here we focus on likewise accelerating the production of liver cells (hepatocytes) from PSCs by first understanding the temporally-dynamic signals controlling liver differentiation.

There has been major headway in generating enriched populations of hepatocyte-like cells from hPSCs (Agarwal et al., 2008; Basma et al., 2009; Cai et al., 2007; Carpentier et al., 2014; Espejel et al., 2010; Han, 2012; Ogawa et al., 2013; Rashid et al., 2010; Si Tayeb et al., 2010; Song et al., 2009; Touboul et al., 2010; Zhao et al., 2012). The considerable interest in generating hepatocytes has stemmed from the fact that they are vital for bodily metabolism and neutralize harmful waste products within the body, amongst other tasks (Stanger, 2015). Attesting to the liver’s importance, acute liver failure rapidly results in coma or death as toxins accumulate in the body (Karl et al., 1953). However the sequence of lineage intermediates through which early hepatocytes develop from pluripotent cells remains to be fully defined, as are the extracellular signals that specify liver fate at each lineage juncture. Importantly, it might be possible to exclusively generate liver cells by deciphering the signals that induce *non-liver* cells at each branching lineage decision and then inhibiting the signals that otherwise specify non-liver fates.

The liver develops through multiple consecutive branching lineage choices, which have been partially delineated through extensive lineage tracing and marker analyses of early vertebrate embryos (Duncan, 2003; Gordillo et al., 2015; Lemaigre, 2009; Lewis and Tam, 2006; Miyajima et al., 2014). In the early mouse embryo, the pluripotent epiblast (at 5.5 days of development [∼E5.5]) differentiates into the anterior primitive streak (∼E6.5) and subsequently, the definitive endoderm germ layer (∼E7-E7.5) (Lawson et al., 1991; Tam and Beddington, 1987). Definitive endoderm is the common progenitor to epithelial cells in diverse internal organs including the liver, pancreas and intestines (Spence et al., 2009; Tremblay and Zaret, 2005). Shortly thereafter by ∼ E8.5, endoderm is patterned along the anterior-posterior axis to broadly form the anterior foregut, posterior foregut and mid/hindgut (Grapin-Botton, 2005; Zorn and Wells, 2009). By ∼E9.5, posterior foregut gives rise to either pancreatic progenitors or the earliest liver progenitors—known as liver bud progenitors (Fukuda-Taira, 1981; Ledouarin, 1964; Rossi et al., 2001)—as shown by single-cell lineage tracing (Chung et al., 2008); conversely, the mid/hindgut gives rise to intestinal epithelium (Spence et al., 2011). Subsequently incipient ∼E9.5 liver bud progenitors are thought to differentiate over the course of several days into either hepatocytes or bile duct cells (cholangiocytes)—the two major epithelial constituents of the liver (Suzuki et al., 2008). Early hepatocytes express characteristic genes (e.g., *Albumin*) (Cascio and Zaret, 1991) but become diversified in fate to form multiple functionally-distinct subsets of hepatocytes after birth (Colnot and Perret, 2011; Jungermann, 1995; Shiojiri et al., 1995; Spijkers et al., 2001), including periportal or pericentral hepatocytes that respectively encircle either portal or central veins (Smith and Campbell, 1988).

Current methods to generate hepatocyte-like cells from hPSCs are often divided into several stages over several weeks or months, generally starting with ACTIVIN/TGFβ (in the presence or absence of WNT) to initially induce endoderm; followed by BMP and FGF to specify liver progenitors; and then HGF, OSM, Dexamethasone and/or 3D reaggregation with other cell-types to specify hepatocytes (Cai et al., 2007; Carpentier et al., 2014; Gouon-Evans et al., 2006; Ogawa et al., 2013; Rashid et al., 2010; Si Tayeb et al., 2010; Song et al., 2009; Takebe et al., 2013; Touboul et al., 2010; Zhao et al., 2012). Each of these stages typically takes several days or weeks, and early HNF4A^+^ liver bud progenitors are typically generated within ∼10-18 days (Ogawa et al., 2013; Si Tayeb et al., 2010; Touboul et al., 2010; Zhao et al., 2012). The use of BMP and FGF to differentiate hPSC-derived endoderm into liver (Cai et al., 2007; Gouon-Evans et al., 2006; Han, 2012; Loh et al., 2014; Si Tayeb et al., 2010; Song et al., 2009; Touboul et al., 2010; Zhao et al., 2012) is rooted in key embryonic explant and genetic studies that indicated the importance of these signals in differentiating endoderm towards liver (Chung et al., 2008; Jung et al., 1999; Rossi et al., 2001; Shin et al., 2007). However both BMP and FGF also specify other endodermal organs including intestines (Loh et al., 2014; Spence et al., 2010). Hence the signals that uniquely induce liver amongst other alternative endodermal lineages remain cryptic.

Here we decompose early liver development into a sequence of six consecutive lineage choices and detail the signals at each juncture that specify each cell-type (either liver or non-liver precursors). We show that multiple developmental signals (e.g., retinoid and other signals) can have opposing effects within 24 hours, initially specifying one fate and then subsequently repressing its formation. This map of liver development allowed us to providently manipulate signals in a temporally-dynamic fashion—and to repress signals at each juncture that specified unwanted cell-types—enabling accelerated production of liver bud progenitors from hPSCs within 6 days. Importantly, the hPSC-derived liver bud progenitors produced in this accelerated fashion are still capable of differentiating into hepatocyte-like cells, the latter of which can engraft the *Fah*^-/-^*Rag2*^-/-^*Ilr2g*^-/-^ mouse model of liver injury. This suggests that accelerated differentiation by precisely manipulating developmental signals does not impede the functionality of downstream cell-types. Finally, to quantitatively track this accelerated liver differentiation process, we furnish new tools; namely cell-surface markers identifying liver progenitors and a knock-in reporter hESC line to track the expression of metabolic enzyme *FAH* during hepatocyte differentiation.

## RESULTS

### RA BMP and FGF activation and TGFβ inhibition rapidly differentiate endoderm into posterior foregut with the competence to later generate liver bud progenitors

Pluripotent cells first differentiate into primitive streak and subsequently definitive endoderm before turning into liver (**Fig. 1a**). We previously identified signals to differentiate hPSCs into >99% pure MIXL1-GFP^+^ primitive streak cells in 24 hours (**Fig. 1b**) and subsequently into >98% pure SOX17-mCherry^+^ definitive endoderm by day 2 of differentiation (**Fig. 1c**) (Loh et al., 2014; Loh et al., 2016). These nearly-pure endoderm populations provided an ideal starting point to examine signals that could further drive day 2 endoderm into day 3 posterior foregut, and later, day 6 liver bud progenitors (**Fig. 1b,c**).

**Figure 1:**
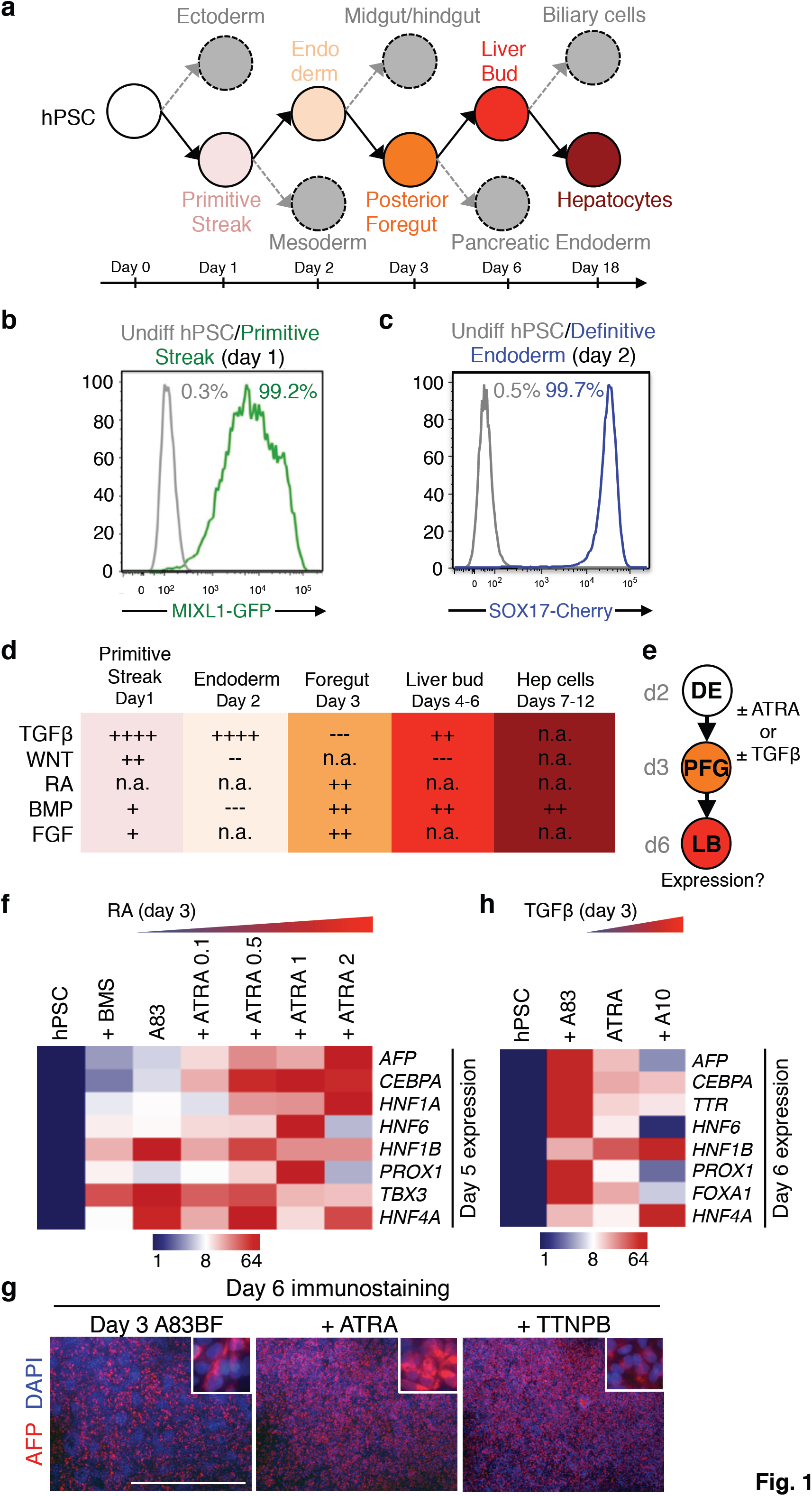
Differentiation of human definitive endoderm into posterior foregut. a) A roadmap for human liver development; summary of the present work. **b**) Percentage of MIXL1-GFP^+^ cells using *MIXL1- GFP* knock-in hESC reporter line. **c**) Percentage of SOX17-mCherry^+^ cells using *SOX17-mCherry* knock-in hESC reporter line. **d**) Temporal dynamism of signals regulating each stage of liver specification (‘+’ = activated; ‘++’ or ‘+++’ = activated at higher levels; ‘-’ = inhibited; ‘n.a.’ = growth factor not added); **e**) Experimental strategy to treat definitive endoderm (DE) with RA or TGFβ modulators on the day 2-3 interval to produce day 3 posterior foregut (PFG) and assaying subsequent effects on liver bud gene expression by day 6, as shown in subpanels **f**,**h**. **f**) qPCR gene expression of day 6 liver bud cells generated from endoderm cells briefly treated on the day 2-3 interval with a retinoid inhibitor (BMS: BMS493, 10μM) or ATRA of varying doses (0.1 μM, 0.5 μM, 1 μM or 2 μM) in the presence of base condition A83 (A83: A8301, 1 μM). **g**) Transient treatment on the day 2-3 interval with ATRA or TTNPB markedly improves AFP expression in day 6 hPSC-derived liver bud progenitors in the presence of base condition A83BF (A83BF: A8301, 1 μM; BMP4, 30 ng/mL; FGF2, 10 ng/mL), as shown by immunostaining with a DAPI nuclear counterstain; scale bar = 1mm. **h**) qPCR gene expression of day 6 liver bud cells generated from endoderm cells briefly treated on the day 2-3 interval with a TGFβ inhibitor A83 (A83: A8301, 1 μM) or a TGFβ agonist (A10: ACTIVIN, 10 ng/mL).

We found that transient activation of the retinoic acid (RA), BMP and FGF pathways together with inhibition of TGFβ signaling for 24 hours was necessary to differentiate day 2 definitive endoderm into day 3 posterior foregut (**Fig. 1d**).

RA pathway agonists initially promoted foregut specification on day 3 of differentiation. Treatment of day 2 endoderm with the RA agonist all-*trans* RA (ATRA, 2 μM) or TTNPB (75 nM) for 24 hours enhanced the formation of day 3 posterior foregut that was later competent to differentiate into liver bud (**Fig. 1e,f,g**) and downstream hepatocytes (**Fig. S1a,b**) on later days of differentiation. Thus RA is crucial for human posterior foregut specification, consistent with how inhibiting RA synthesis abrogates both liver and pancreas formation in zebrafish embryos (Stafford and Prince, 2002).

Foregut specification by day 3 was also promoted by TGFβ inhibition (**Fig. 1h**) and activation of the BMP and FGF pathways (**Fig. S1d,e,h,i**). Each of these manipulations enhanced the generation of posterior foregut that was capable of subsequently differentiating into liver bud progenitors and/or hepatocytes (**Fig. 1h**; **Fig. S1c**). Taken together, day 2 definitive endoderm could be converted into day 3 posterior foregut by the simultaneous activation of RA, BMP and FGF pathways together with inhibition of TGFβ signaling for 24 hours; such day 3 posterior foregut populations expressed *HHEX* (**Fig. S1j**), a marker of ∼E8.5 mouse ventral posterior foregut (Thomas et al., 1998).

### TGFβ, BMP, PKA activation and WNT inhibition accelerate differentiation of foregut into liver bud progenitors by day 6 of differentiation

Though day 3 posterior foregut was initially specified by RA activation and TGFβ inhibition, the subsequent differentiation of posterior foregut into day 6 liver bud progenitors was *suppressed* by continued RA activation and TGFβ inhibition. Instead, liver bud specification on days 4-6 required activation of the TGFβ, BMP and PKA pathways, together with WNT inhibition, for 72 hours (**Fig. 2a**).

**Figure 2:**
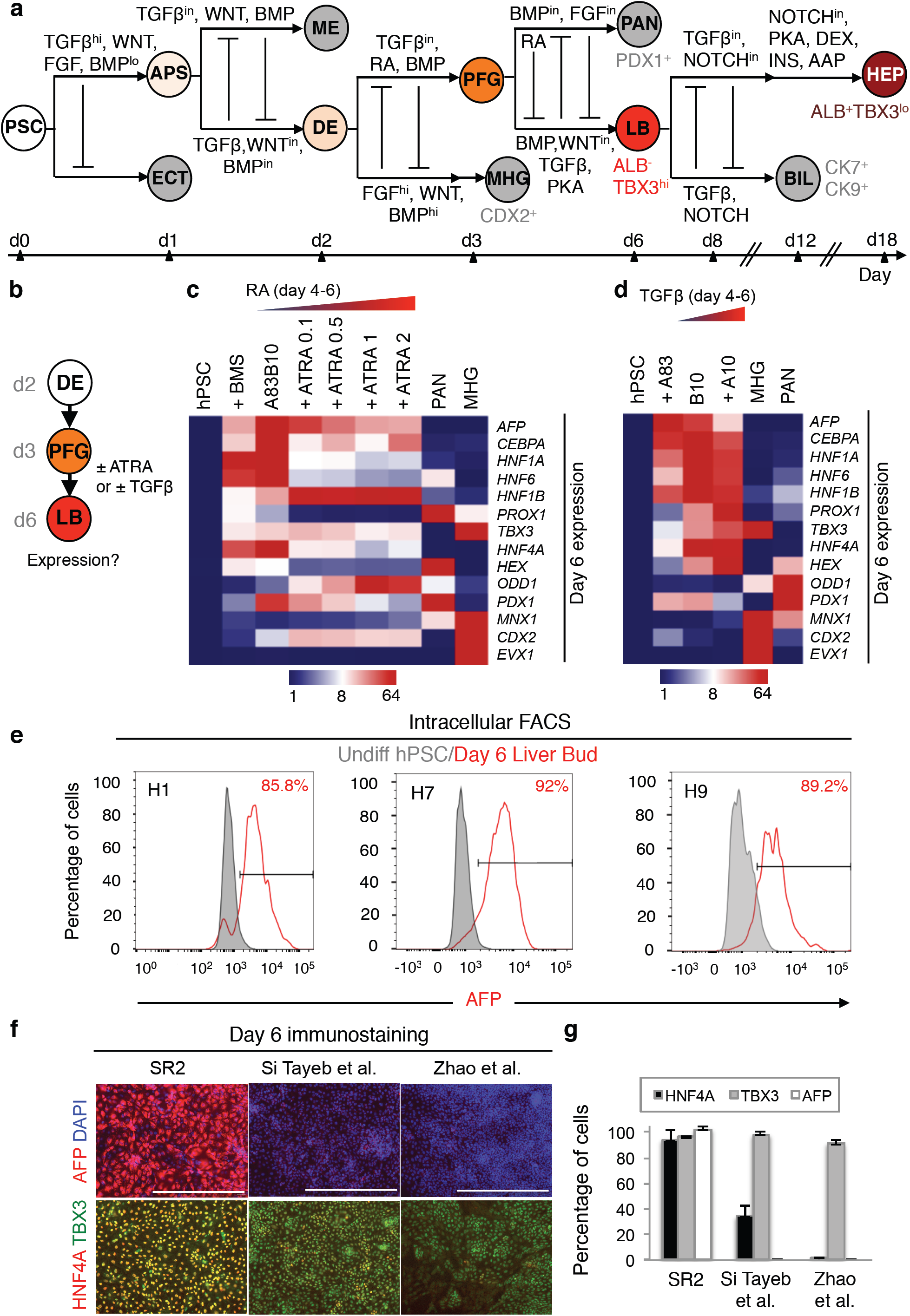
Accelerated generation of human TBX3^+^HNF4A^+^AFP^+^ liver bud progenitors by day 6 of PSC differentiation. **a)** A signaling roadmap for human liver development; summary of the present work: anterior primitive streak (APS), definitive endoderm (DE), posterior foregut (PFG), liver bud (LB), pancreatic endoderm (PAN), cholangiocytes/biliary (BIL) cells, immature hepatocyte-like cells (HEP), low signals (lo), high signals (hi), signaling inhibitor (in). **b**) Experimental strategy to treat posterior foregut (PFG) with RA or TGFβ modulators on the day 4-6 interval to produce day 6 liver bud progenitors (LB) and assaying effects on liver bud gene expression by day 6, as shown in subpanels **c**-**d**. **c**) qPCR gene expression of day 6 liver bud cells generated from endoderm treated on the day 4-6 interval with a retinoid inhibitor (BMS: BMS493, 10 μM) or varying doses of a retinoid agonist (ATRA, 0.1 μM, 0.5 μM, 1 μM, and 2 μM) in the presence of base condition A83B10 (A83B10: A8301, 1 μM; BMP4, 10 ng/mL) and qPCR gene expression of day 6 hPSC-derived mid/hindgut (MHG) or pancreatic endoderm (PAN) cells. **d**) qPCR gene expression of day 6 liver bud cells generated from endoderm treated on the day 4-6 interval with a TGFβ inhibitor (A83: A8301, 1 μM) or ACTIVIN (10 ng/mL) in the presence of base condition B10 (B10: BMP4, 10 ng/mL). Day 6 hPSC-derived pancreatic endoderm (PAN) or mid/hindgut cells (MHG) were included as controls. **e**) Percentage of day 6 liver bud progenitors positive for AFP from the differentiation of 3 hPSC lines (H1, H7 and H9) as shown by intracellular FACS. **f**) Day 6 H1 hPSC-derived liver progenitors generated by 3 methods: SR2 (the method described in the present work) or previously-reported methods (Si Tayeb et al., 2010; Zhao et al., 2012) were immunostained for AFP, HNF4A and TBX3 with a DAPI nuclear counterstain; scale bar = 500μm. **g**) Percentages of AFP^+^, TBX3^+^ or HNF4A^+^ day 6 liver bud progenitors counted using ImageJ analyses of images in subpanel **f**.

While RA initially differentiated endoderm into day 3 posterior foregut (above), 24 hours later it repressed subsequent progression into liver bud progenitors on days 4-6 (**Fig. 2b,c**). Hence our findings reconcile conflicting findings that RA is overall required for zebrafish liver induction (Stafford and Prince, 2002) but that RA-coupled beads inhibit liver bud marker formation in HH10 stage zebrafish embryos (Bayha et al., 2009)—there is a temporally dynamic requirement for RA in liver bud specification.

Akin to RA signaling, the role for TGFβ in liver specification was also temporally dynamic: TGFβ inhibition initially promoted foregut formation by day 3 (**Fig. 1e,h**) but 24 hours later, TGFβ *activation* promoted liver bud specification on days 4-6, leading to enhanced liver bud marker expression by day 6 (**Fig. 2b,d**). Emphasizing the importance of TGFβ activation, we found that TGFβ inhibition at this stage instead abrogated the differentiation of foregut into liver bud (**Fig. 2b,d**), contrasting with earlier use of TGFβ inhibitors to differentiate hPSC-derived endoderm into liver (Loh et al., 2014; Sampaziotis et al., 2015; Touboul et al., 2010).

BMP and PKA activation, together with WNT blockade, also cooperated with TGFβ activation to differentiate day 3 foregut into day 6 liver bud progenitors. Consistent with earlier findings (Chung et al., 2008; Rossi et al., 2001; Shin et al., 2007; Si Tayeb et al., 2010; Wandzioch and Zaret, 2009; Zhao et al., 2012), first we found that BMP activation differentiated foregut into liver bud while blocking pancreas formation; by contrast BMP inhibition suppressed liver formation (**Fig. S2a**,**g**). This mirrors how *bmp2b* overexpression promotes liver specification at the expense of pancreatic progenitors in zebrafish embryos (Chung et al., 2008) and how BMP induces liver from mouse embryonic endoderm explants (Rossi et al., 2001). Second, we found PKA agonists (e.g., 8-bromo-cAMP) also potently specified liver bud from foregut (**Fig. S2b**), supporting the notion that prostaglandin E2 specifies zebrafish liver progenitors by activating the PKA cascade (Nissim et al., 2014). Finally, WNT inhibition (using C59) from days 4 to 6 suppressed mid/hindgut (MHG; posterior endoderm) formation (**Fig. S1f-g**) and instead promoted the formation of liver bud progenitors that had the ability to subsequently form hepatocytes (**Fig. S2f**) and cholangiocytes (**Fig. S4g**). This is consistent with how WNT signaling blocks foregut formation and instead specifies MHG in *Xenopus* (McLin et al., 2007). Taken together, simultaneous activation of TGFβ, BMP and PKA but suppression of WNT drove day 3 posterior foregut into day 6 liver bud progenitors while simultaneously blocking pancreatic and MHG differentiation.

Combining the above signals enabled the generation of a rather-uniform liver bud progenitor population by day 6 of hPSC differentiation, which was accelerated by comparison to extant methods. Quantification of differentiation efficiencies by intracellular flow cytometry demonstrated the consistent generation of an 89.0±3.1% pure AFP^+^ liver bud progenitor population from the H1, H7 and H9 hPSC lines (**Fig. 2e**). This new combined approach (“SR2”) for liver bud specification was more rapid than extant liver induction methods (Si Tayeb et al., 2010; Zhao et al., 2012), yielding significantly higher expression of liver bud markers and purer populations of liver bud progenitors (94.1±7.35% HNF4A^+^) by day 6 of hPSC differentiation (**Fig. 2e,f**,**g**; **Fig. S2i**), whereas HNF4A^+^ liver bud progenitors were instead efficiently formed on later days (by day 14) in other differentiation protocols (Si Tayeb et al., 2010; Zhao et al., 2012).

Finally day 6 hPSC-derived liver bud progenitors expressed high levels of liver bud transcription factors including *AFP*, *HNF4A*, *TBX3*, *HNF6* and *CEBPA*, which were low or undetectable in MHG (**Fig. S2c-e**). Reciprocally hPSC-derived liver bud progenitors did not express *CDX* or *HOX* genes (**Fig. S2c-e**), which are markers of the MHG, a developmentally-related lineage that emanates from adjacent posterior endoderm and is also specified by BMP and FGF signals (Sherwood et al., 2011; Spence et al., 2010).

Taken together, though liver bud and MHG (intestinal) progenitors are spatially adjacent *in vivo*, they are transcriptionally distinct lineages and can be produced in mutually-exclusive signaling conditions *in vitro* from hPSCs. Importantly, while there is a common requirement for BMP and FGF in liver and MHG specification, we reveal signals that uniquely specify liver (**Fig. 2a)**, thus clarifying how these lineages become segregated from one another. This knowledge enabled the efficient generation of human liver bud progenitors by day 6 of PSC differentiation, which is >2 times faster than extant methods.

### A surface marker signature for hPSC-derived liver bud progenitors

To quantitatively track this accelerated time course of human liver bud progenitor specification at the single-cell level, we next sought to identify cell-surface markers that would demarcate day 6 liver bud progenitors from developmentally-earlier fates (day 0 hPSCs and day 2 definitive endoderm). Systematically screening the expression of 242 cell-surface antigens on these 3 lineages revealed stage-specific surface markers (**Fig. 3a,b**). First, CD10 was expressed in 92.6±5.6% of undifferentiated hPSCs but was abruptly downregulated upon differentiation, being expressed in <2% of cells in day 2 definitive endoderm or day 6 liver bud populations (**Fig. 3c**; **Fig. S3b**; **Table S1**). Second, CD184/CXCR4 (a known definitive endoderm marker (D’amour et al., 2005)) was enriched in day 2 definitive endoderm by comparison to day 0 hPSCs and day 6 liver bud progenitors (**Fig. 3c,d; Table S1**). Finally, CD99 was highly expressed on day 6 liver bud progenitors by comparison to preceding hPSCs or DE (**Fig. 3c,d; Table S1**). This pattern was consistent across a panel of 4 hPSC lines (H7, HES2, H1 and BJC1; **Table S1**; **Fig. S3a,b**). Thus, positive selection for CD99^hi^ cells may enrich for liver bud progenitors, especially when combined with negative selection to eliminate CD10^+^ hPSCs and CD184^+^ DE. These cell-surface markers can be used to track early liver progenitor specification and to distinguish them from developmentally-earlier cell-types.

**Figure 3:**
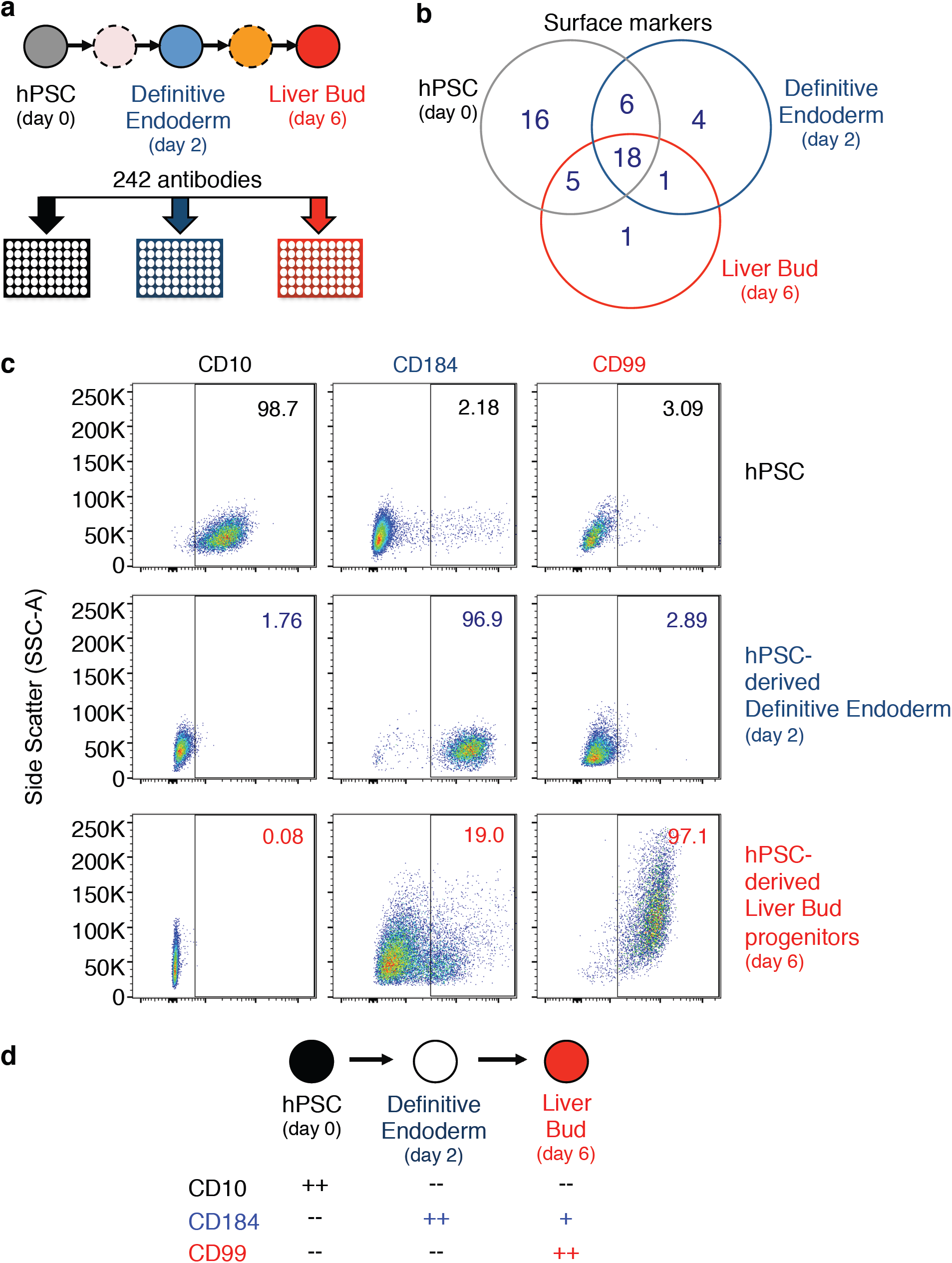
High-throughput screening identifies liver-specific surface markers during PSC differentiation. **a)** Experimental strategy to conduct high-throughput FACS screening of surface markers expressed on H7 hPSCs, day 2 H7 hPSC-derived endoderm and day 6 H7 hPSC-derived liver bud progenitors. **b**) Venn diagram of surface markers expressed on hPSCs, day 2 hPSC-derived DE and day 6 hPSC-derived liver bud progenitors. **c**) hPSCs, hPSC-derived definitive endoderm and liver bud progenitors stained for CD10, CD184 and CD99 as shown by live-cell FACS. **d**) Surface markers expressed on hPSC and day 2 hPSC-derived definitive endoderm and day 6 liver bud progenitors; summary of present work.

### Segregation of human liver bud progenitors into hepatocyte vs. biliary fates by competing NOTCH, TGFβ, PKA and other signals

Next we tested whether these hPSC-derived day 6 liver bud progenitors—obtained via accelerated differentiation—were still competent to subsequently give rise to *ALBUMIN*^+^ hepatocyte-like cells or *SOX9*^+^ biliary cells (cholangiocytes) on later days of differentiation. Suppression of both NOTCH and TGFβ signaling on days 7 to 8 was necessary to block biliary formation and consolidate hepatocyte fate. First, we found that NOTCH blockade (by DAPT) prevented hPSC-derived liver bud progenitors from adopting a biliary fate (as assessed by reduced S*OX9* expression) and instead diverted cells into *ALBUMIN*^+^ hepatocytes (**Fig**. **S4a,b,e**). Second, brief TGFβ blockade after the liver bud stage also enhanced expression of various liver genes including *ALBUMIN* (**Fig. S4c,d**). Reciprocally, TGFβ activation (using ACTIVIN) reduced *ALBUMIN* expression but induced *SOX9* expression in a dose-dependent fashion (**Fig. S4f**). Collectively, NOTCH and TGFβ signaling drive biliary differentiation and hence must be suppressed to promote hepatocyte fate from liver bud progenitors.

Conversely, consistent with earlier studies (Ogawa et al., 2015; Sampaziotis et al., 2015), activation of TGFβ and NOTCH together with insulin differentiated day 6 liver bud progenitors into CK7^+^/CK19^+^ biliary progenitors by day 13 of hPSC differentiation (**Fig. S4g**). This suggests the bipotent capacity of liver bud progenitors at a population level to give rise to either hepatic or biliary fates (which remains to be formally demonstrated *in vivo* by single-cell lineage tracing) and identifies the signals that control the mutually-exclusive allocation of hepatocyte-like cells vs. cholangiocytes.

Beyond expression of pan-hepatocyte marker *ALBUMIN*, we sought to assess what other key liver enzymes and markers were expressed by hPSC-derived hepatocyte-like cells and we screened for signals that could augment the acquisition of certain physiologic characteristics. In particular we sought to promote expression of tyrosine metabolism pathway components in hPSC-derived hepatocytes, as genetic deficiency of tyrosine metabolic enzymes (e.g., *fumarylacetoacetate hydrolase* [*FAH*]) results in hereditary tyrosinemia in human patients, a metabolic disorder (St-Louis and Tanguay, 1997). High insulin levels together with a stabilized ascorbic acid derivative (ascorbic acid-2-phosphate [AAP]) greatly promoted the expression of tyrosine metabolic pathway genes *PAH*, *HGD*, *HPD*, *TAT*, *MAI* and *FAH* and other liver markers when applied on days 7-12 of hPSC differentiation (**Fig. 4b, c**). PKA agonists (e.g., 8-bromo-cAMP and forskolin) also had a similar effect (**Fig. S4h**). Microarray profiling showed treatment with NOTCH inhibitors, AAP, forskolin or insulin upregulated genes associated with gene ontology classifications pertaining to metabolic processes (**Fig. 4a**; **Fig. S4i**; **Table S5**), asserting that these individual manipulations each promote the metabolic competence of hPSC- derived hepatocytes.

**Figure 4:**
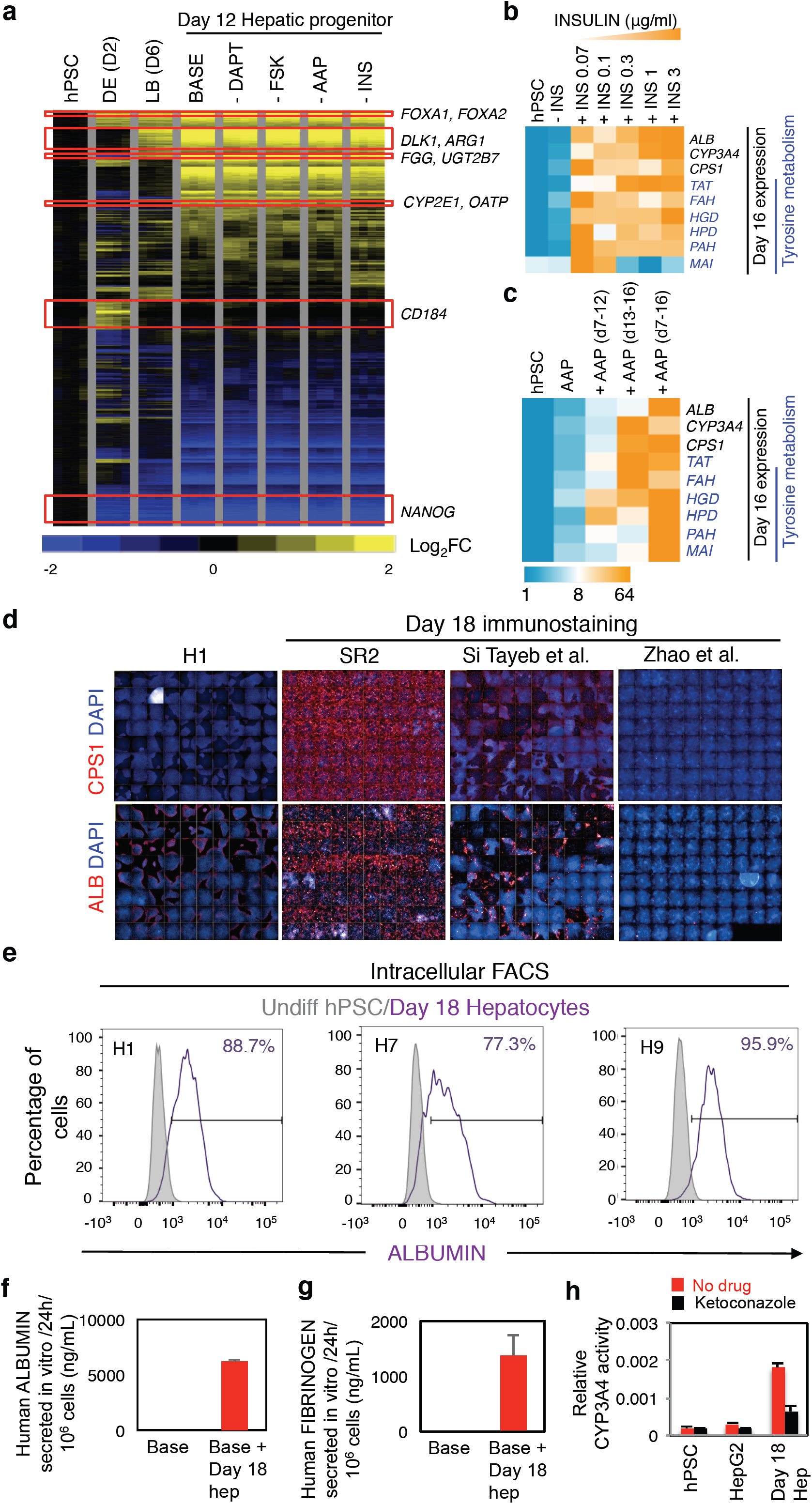
Generation of enriched populations of ALB^+^ and FAH^+^ immature hepatocytes by day 18 of PSC differentiation. **a)** Global microarray gene expression of hPSCs, day 2 DE, day 6 liver bud (LB) progenitors and day 12 hPSC-derived hepatic progenitors that were either induced using the full combination (base) or with the individual omission of either the NOTCH inhibitor (DAPT), forskolin (FSK), ascorbic acid-2-phosphate (AAP) or insulin (INS) during the liver bud →→ hepatic progenitor differentiation step to reveal genes whose expression is regulated by DAPT, FSK, AAP and INS. 4 biological replicates were used for the microarray analyses. **b**) qPCR gene expression of day 16 hepatocytes generated from liver bud progenitors treated on the day 7-12 interval with INSULIN (INS: 0.7, 1, 3, 10 and 30 μg/mL). **c)** qPCR gene expression of day 16 hepatocytes generated from liver bud progenitors treated on the day 7-12, 12-16 or 7-16 intervals with <u>a</u>scorbic <u>a</u>cid 2-<u>p</u>hosphate (AAP, 200 μg/mL). **d**) Day 18 hepatocyte-like cells generated by 3 methods: SR2 (the method described in the present work) or previously-reported methods (Si Tayeb et al., 2010; Zhao et al., 2012) were immunostained for Carbamoyl Phosphate Synthetase 1 (CPS1) and ALBUMIN (ALB) with a DAPI nuclear counterstain and multiple fields were stitched into 1 image. **e**) ALBUMIN intracellular FACS analysis of hPSCs or day 18 hepatocytes derived from hPSC lines H1, H7 and H9. **f**) ALBUMIN levels detected in culture medium grown with hPSC-derived hepatocytes as measured by ELISA with a human-specific ALBUMIN antibody; “Base” denotes the culture medium alone. **g**) FIBRINOGEN levels detected in culture medium grown with hPSC-derived hepatocytes as measured by ELISA; “Base” denotes the culture medium alone. **h**) CYP3A4 activity of hPSCs, HepG2 and hPSC-derived hepatocytes (Day 18 Hep) as measured by P450-Glo luciferase assays and abolished by CYP3A4 inhibitor ketoconazole; signal is further normalized to cell number baseline as measured by Cell Titer Glo.

Coordinated application of these hepatocyte-specifying signals—together with inhibition of cholangiocyte-specifying signals TGFβ and NOTCH—enabled the generation of an enriched population of ALB^+^ hepatocyte-like cells by day 18 of hPSC differentiation (**Fig. 4d; Fig. S4j)**. Intracellular flow cytometric analysis revealed the consistent generation of, on average, an 87.3±9.4% pure ALBUMIN^+^ hepatocyte population from the H1, H7 and H9 hPSC lines (**Fig. 4e**). This new combined approach yielded significantly higher expression of hepatocyte markers such as ALBUMIN and CPS1 within a span of 18 days when compared with other methods (Si Tayeb et al., 2010; Zhao et al., 2012) at the same timepoint, as revealed by widefield fluorescent imaging of entire wells of differentiated cells (**Fig. 4d**).

We further tested if the coordinated application of these signals enabled the generation of hepatocyte like-cells that secreted ALBUMIN and possessed P450 cytochrome activities, which other differentiation methods have successfully attained (Basma et al., 2009; Carpentier et al., 2014; Si Tayeb et al., 2010; Zhao et al., 2012). First, day 18 hPSC-derived hepatocytes stained positive for periodic acid Schiff (**Fig. S4k**). Second, they secreted 5.89±0.71 μg/mL of ALBUMIN protein (**Fig. 4f**) and 1.38±0.26 μg/mL of FIBRINOGEN protein into the medium (**Fig. 4g**)—levels in the same order of magnitude as the amount secreted *in vivo* by hepatocytes in human beings (Bernardi et al., 2012; Tennent et al., 2007). Third, they also possessed CYP3A4 enzymatic activity at levels higher than that in HepG2 cells, which was attenuated by the CYP3A4 inhibitor ketoconazole (**Fig. 4h**). Given that these three criteria have been previously applied to characterize hPSC-derived hepatocyte-like cells (Espejel et al., 2010; Ogawa et al., 2013; Rashid et al., 2010; Si Tayeb et al., 2010; Takebe et al., 2013; Zhao et al., 2012), we sought to further focus on expression of tyrosine metabolic pathway enzymes (**Fig. 4b,c; Fig. S4a-d**), which are pertinent to the treatment of the *Fah*^-/-^*Rag2*^-/-^*Ilr2g*^-/-^ mouse model of hereditary tyrosinemia (described below).

To track the expression of FAH at a single-cell level in hPSC-derived hepatocytes, we used CRISPR/Cas9-mediated gene editing to generate a *FAH-2A-Clover* knock-in H1 hESC reporter line in which the FAH coding sequence was left intact (**Fig. 5a,b**). Optimal guide RNAs that efficiently directed Cas9 cleavage of the FAH locus were first evaluated in HEK293T cells (**Fig. S5a-c**) and then used to direct homologous recombination in hESCs to knock-in a *Clover* fluorescent reporter gene downstream of *FAH*. Allele-specific integration of the reporter cassette was confirmed by PCR (**Fig. S5d-f**) and sequencing (**Fig. S5g**). Applying the above differentiation protocol to the resultant *FAH-2A-Clover* hESCs yielded 81.5±3.2% FAH-Clover^+^ population in the day 18 hepatocyte-like population (**Fig. 5c,d**). Attesting to the faithfulness of the reporter, FACS-sorted Clover^+^ day 18 cells were significantly enriched for *FAH*, *ALBUMIN* and *HGD* mRNAs by comparison to Clover^-^ cells (**Fig. 5e,f**). The expression of FAH in the majority of hPSC-derived hepatocyte-like cells motivated us to test whether these populations could be used to treat the *Fah*^-/-^*Rag2*^-/-^*Ilr2g*^-/-^ mouse model of liver injury.

**Figure 5:**
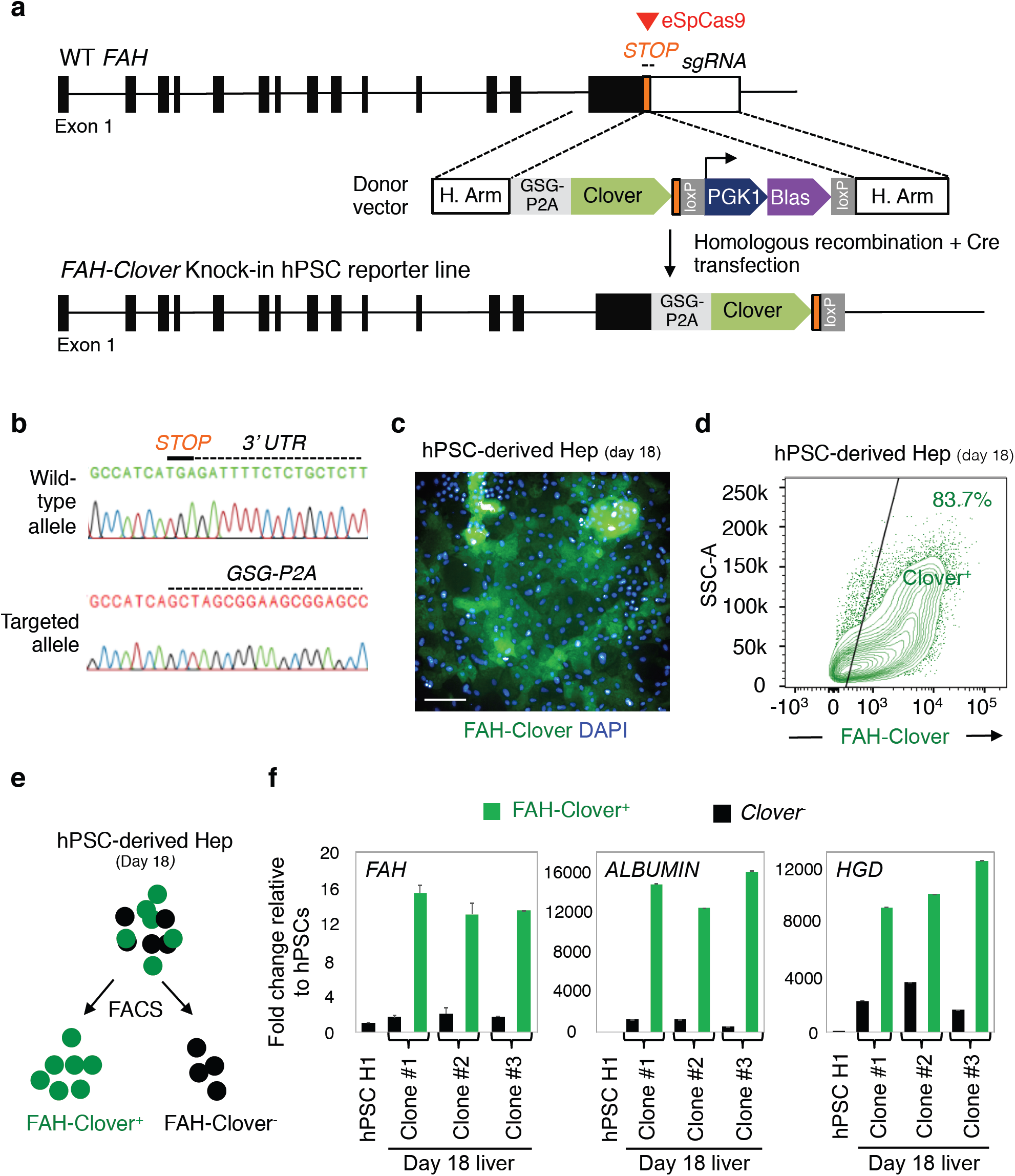
Generation of a *FAH-2A-Clover* knock-in H1 human pluripotent stem cell (hPSC) reporter line using CRISPR/Cas9-mediated gene editing. **a)** Schematic illustration of the strategy to insert a P2A- Clover reporter cassette into the *FAH* locus. A Gly-Ser-Gly (gsg) sequence was added to the start of the P2A sequence; ‘H. Arm’ = homology arms; ‘eSpCas9’ = enhanced specificity Cas9; ‘PGK1’ = Phosphoglycerate kinase 1 promoter, ‘Blas’ = Blasticidin drug resistance gene. **b**) Wild-type and knock-in alleles of the same locus sequenced by PCR. **c**) Immunofluorescence of FAH-Clover^+^ cells co-stained with DAPI, scale bar = 100μm. **d**) 83.7% FAH-Clover^+^ population in day 18 hPSC-derived hepatocyte-like population (hPSC-derived Hep). **e**) Schematic illustration showing the separation of FAH-Clover^+^ and FAH- Clover^-^ cell fractions within d18 hPSC-derived hepatocyte-like cells by fluorescence activated cell sorting (FACS). **f**) FACS-sorted FAH-Clover^+^ day 18 H1 derived hepatocyte-like cells (green bars) were significantly enriched for liver genes *FAH, ALBUMIN* and *HGD* mRNAs when compared with FAH-Clover^-^ cells (black bars). Consistent results in various *FAH-2A-Clover* knock-in H1 hPSC reporter clones #1-3.

### hPSC-derived hepatocytes engraft the injured mouse liver and improve survival during liver injury

Finally we sought to determine whether bulk populations of hPSC-derived day 18 hepatocyte-like cells generated using the above signaling strategy could engraft the injured mouse liver and improve short-term survival. Specifically, we tested whether *FAH*^+^ hPSC-derived hepatocytes derived using our differentiation schema could engraft a genetic mouse model of liver injury— namely *Fah*^-/-^*Rag2*^-/-^*Il2rg*^-/-^ (FRG) mice (Azuma et al., 2007; Overturf et al., 1996). FRG mice are an immunodeficient mouse model of Tyrosinemia Type I, an inherited liver failure syndrome caused by *FAH* mutations in patients (Labelle et al., 1993; St-Louis and Tanguay, 1997). Like human patients, FRG mice suffer severe and eventually fatal liver injury in the absence of a hepatoprotective drug, NTBC (Azuma et al., 2007; Overturf et al., 1996).

To this end, day 18 hPSC-derived hepatocytes were doubly labeled with constitutively-expressed genetic reporters (*EF1A-BCL2-2A-GFP* and *UBC-tdTomato-2A-Luciferase*) (Loh et al., 2016) and then were intrahepatically injected into neonatal FRG mice (less than 2 days old). Injected mice were allowed to grow until 6 weeks of age before liver injury was induced by cyclical withdrawal of hepatoprotective drug NTBC (Azuma et al., 2007; Overturf et al., 1996; Zhu et al., 2014) (**Fig. 6a**). Strikingly, hPSC-derived hepatocytes engrafted in 7 out of 15 mice (**Fig. 6b**) and continued to expand *in vivo* as shown by an increase in bioluminescence intensity over time (**Fig. 6b, S6a**). After ∼10 weeks of liver injury, all surviving mice had detectable bioluminescence (indicating the presence of transplanted cells); mice that initially had little/no bioluminescence had died by this timepoint (**Fig. 6b**). This suggests that engrafted hPSC-derived hepatocytes promoted the survival of the FRG mice. Bioluminescent imaging of dissected livers (**Fig. S6b,c**) and sectioning revealed the presence of human ALBUMIN^+^ tdTomato^+^ liver cells near the vasculature within the livers of mice that survived 5 months post-transplantation (**Fig. S6b,c**).

**Figure 6:**
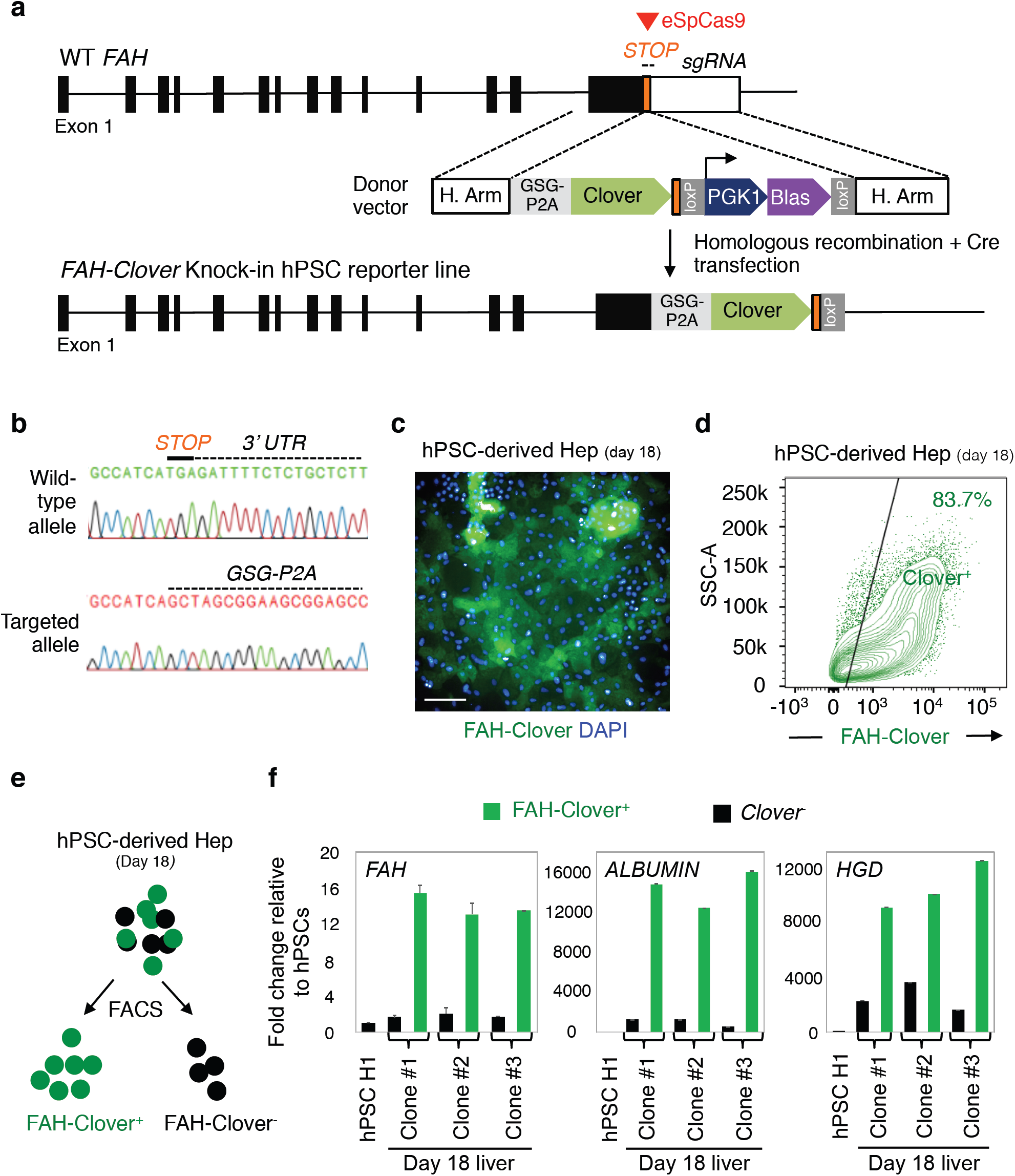
Human PSC-derived hepatocytes engraft neonatal and adult *Fah*^-/-^*Rag2*^-/-^*Ilr2g*^-/-^ mice and improve short-term survival. **a)** Experimental strategy to inject hPSC-derived hepatocytes into the livers of neonatal FRG mice (<48 hrs old) or adult FRG mice (4-6 weeks old) and to subsequently induce liver injury; H9 *EF1A-BCL2-2A-IRES*-*UBC-tdTomato-2A-Luciferase* hPSCs were used to generate hepatocytes for injections. (**b**) Survival records of adult FRG mice that have been intrahepatically injected with Luciferase^+^ day 18 hPSC-derived hepatocytes while they were neonates. Liver injury was induced by withdrawing NTBC for 3 or 10 weeks. **c**) Luciferase^+^ day 18 hPSC-derived hepatocytes engrafted in adult FRG mice. **d**) Day 18 hPSC-derived hepatocytes engrafted in mice liver and stained for human ALBUMIN (brown)(arrowheads) at various magnifications as shown by immunostaining; scale bar = 200μm; H9 *EF1A- BCL2-2A-GFP* were used. **e**) Kaplan-Meier’s survival curve depicting the percentage survival of mice that had either been injected with ∼1.5 million day 18 hPSC-derived hepatocytes (red line) (*N*=11) or media-only control (blue line) (*N*=12); Mantel-Cox log-rank test, *p-value<0.05, result from 3 independent experiments. **f**) Reduction of bilirubin levels in serum in mice transplanted with hPSC-derived hepatocytes vs. negative or no-cell media only control. **g**) Human serum ALBUMIN secretion into blood stream in vivo by hPSC-derived hepatocytes as measured by ELISA.

We further tested the ability of our hPSC-derived hepatocytes to ameliorate liver failure in a second model by intrasplenically transplanting them into 4- to 6-week-old *adult* FRG mice (**Fig. S6d**) and inducing chronic liver injury by intermittent NTBC withdrawal (**Fig. 6a**). One month post-transplantation, human ALBUMIN^+^ liver cells were detected in the mouse liver, some of which were localized near the vasculature (**Fig. 6d**; **Fig. S6e**). On average 120±36 ng/mL of human ALBUMIN was detected in the mouse bloodstream (*N* = 7 mice) indicating modest ALBUMIN secretion by the transplanted hPSC-derived hepatocytes (**Fig. 6g**). Bilirubin levels (reflecting the extent of liver injury) were reduced in the serum of these mice (**Fig. 6f**). 72.7% of the FRG mice transplanted with hPSC-derived hepatocytes survived longer compared to the negative control mice that were not injected with any cells (**Fig. 6e**). Together, these results indicated that transplanted hPSC-derived hepatocytes could engraft the injured adult mice liver, secrete human serum ALBUMIN and ameliorate liver injury. This demonstrates that hPSC-derived liver bud progenitors obtained via accelerated differentiation are still competent to generate hepatocyte-like cells capable of some degree of *in vivo* engraftment and that they can improve short-term survival in the FRG mouse model of hereditary tyrosinemia.

## DISCUSSION

Contrary to the view that hPSC differentiation into various cell-types is innately a protracted process, multiple groups including our own have shown that precisely timed manipulation of extracellular signals can accelerate differentiation (Chambers et al., 2012; Loh et al., 2016; Qi et al., 2017). Indeed during development, differentiating progenitors interpret signals in a temporally-dynamic way, with drastically different responses in even a 24-hour interval (Wandzioch and Zaret, 2009). Thus manipulating these signals *in vitro* with equal dynamism is necessary to achieve efficient differentiation. Here we show that starting from highly-pure definitive endoderm populations, it is possible to generate enriched liver bud progenitors by day 6 of hPSC differentiation while suppressing differentiation into alternate endodermal fates (pancreas and MHG). These liver bud progenitors are produced >2 times faster than possible by extant methods. Though derived in accelerated fashion, these hPSC-derived liver bud progenitors are still competent to differentiate into hepatocyte-like cells and cholangiocytes at later days. Through the use of *FAH-2A-Clover* reporter hESCs, we demonstrate that the resultant hepatocyte-like cells express tyrosine metabolic enzyme FAH and that accordingly, they can engraft and improve short-term survival in the *Fah*^-/-^*Rag2*^-/-^*Il2rg*^-/-^ mouse model of hereditary tyrosinemia. It is hoped that this accelerated system to generate hPSC-derived liver bud progenitors will avail the final goal of generating mature hepatocytes, which has yet to be realized.

Our work highlighted two principles to more specifically generate liver at the expense of other endodermal lineages. First, we found that signals that specified liver fate were temporally dynamic, initially promoting but then inhibiting liver fate. Indeed over a brief 24-hour interval, RA activation and TGFβ blockade were necessary to drive definitive endoderm into posterior foregut progenitors, but the next day, these same signals blocked the further progression of posterior foregut towards a liver bud fate. Hence understanding the temporal dynamics with which these signals act is key, as these signals must be manipulated with equal dynamism to guide efficient differentiation as cells pass through transient windows of inductive competence.

Second, at each lineage segregation, we determined the signals that promoted the formation of both our desired fate as well as the unwanted lineage(s). With this knowledge, these lineage segregations could be efficiently negotiated by providing the relevant *inductive* signal(s) to drive differentiation towards the desired lineage while *repressing* the signal(s) that otherwise promoted the alternate fate (Loh et al., 2014; Loh et al., 2016). For example, NOTCH and TGFβ signaling differentiated liver bud progenitors into cholangiocytes while suppressing hepatocyte formation. Conversely, blockade of NOTCH and TGFβ pathways induced differentiation into hepatocytes, in part by excluding formation of SOX9^+^ cholangiocytes.

Taken in collective our results suggest a signaling code for accelerated liver specification (**Fig. 2a**), extending beyond the use of BMP and FGF to induce liver progenitors from endoderm (Gouon-Evans et al., 2006; Shiraki et al., 2008; Si Tayeb et al., 2010; Spence et al., 2010; Touboul et al., 2010; Zhao et al., 2012). Indeed, liver and MHG both share a common requirement for BMP and FGF, raising the question of how these two lineages are distinguished from one another. We show that transient exposure of endoderm to RA activation and TGFβ blockade specifies the competence of posterior foregut, such that upon subsequent exposure to BMP and FGF, it specifically differentiates into liver (instead of MHG). Moreover, while we confirm that BMP and FGF are necessary to differentiate posterior foregut into liver bud, this is only fully realized in the context of other signals (TGFβ and PKA activation together with WNT inhibition). We also introduce new tools to track this accelerated differentiation process: (i) the surface marker CD99 which, together with absence of CD10 and CD184/CXCR4, identifies day 6 liver bud progenitors and (ii) an *FAH-Clover* reporter hESC line to track the subsequent generation of *FAH*^+^ hepatocyte-like cells.

Finally, we show that hPSC-derived hepatocyte-like cells, though derived in accelerated fashion, are competent to engraft *in vivo* and improve the survival of *Fah*^-/-^*Rag2*^-/-^*Il2rg*^-/-^ mice. Substantial progress has been made in generating human hepatocytes from direct lineage reprogramming that can engraft sublethal models of mouse liver injury (Huang et al., 2014; Sekiya and Suzuki, 2011; Song et al., 2016; Yu et al., 2013; Zhu et al., 2014). However, generating functionally mature human hepatocytes from hPSCs remains a significant challenge. It is hoped that human liver differentiation roadmap described here and this accelerated system to generate liver bud progenitors will provide insight into human liver development and avail the as-of-yet-unrealized goal of generating mature hepatocytes.

## EXPERIMENTAL PROCEDURES

### hPSC differentiation and hepatocyte culture

mTeSR1-grown hPSCs (H1, H7 and H9 from WiCell and ESI035 from ESI BIO) were tested to be mycoplasma-free and were seeded for differentiation as single cells using Accutase (Millipore, SCR005) onto Geltrex-coated plates (Thermo Fisher, A1413302) at a density of 40,000 cells per well of 12-well plate. Definitive endoderm (DE) was generated from hPSCs through the anterior primitive streak intermediate using DE Induction Medium A for 24 hours, followed by DE Induction Medium B for 24 hours (Thermo Fisher, A27654SA) (Loh et al., 2014; Loh et al., 2016). DE was then patterned into posterior foregut (PFG) by using A8301 (1 μM) and ATRA (2 μM) in CDM3 medium for another 24 hrs. PFG was further differentiated to liver bud progenitors using ACTIVIN (10 ng/mL), 8-bromo-cAMP (1 mM), C59 (1 μM) and BMP4 (30 ng/mL) in CDM3 medium for 3 days, followed by BMP4 (10 ng/mL), Oncostatin M (10 ng/mL), Dexamethasone (10 μM), RO4929097 (2 μM) or DAPT (10 μM), forskolin (10 μM), human recombinant insulin (10 μg/mL), ascorbic acid-2 phosphate (AAP; 200 μg/mL) and amino acid concentrate in CDM4 base medium for the next 6 days (day 7 to 12). SB505124 (1μM) was added for 48 hours on days 7 and 8. Subsequently, hPSC-derived hepatocytes were further treated with Dexamethasone (10 μM), RO4929097 (2 μM) or DAPT (10 μM), forskolin (10 μM), human recombinant insulin (10 μg/mL) and AAP (200 μg/mL) in CDM5 base medium for 6 days (day 13 to 18). Cryopreserved adult human hepatocytes (Thermofisher Scientific, HU1352, HU8055 or HU8132) were thawed with CHRM (Gibco, CM7000), plated and grown in sandwich cultures (Gibco, CM3000, CM4000) as per manufacturer’s instructions. For detailed methods, see **Supplementary Information**.

### Animal handling and transplantation

FRG^-/-^ C57BL6 or FRG^-/-^ NOD (Yecuris) mice were fed *ab libitum* with low-protein high-fat irradiated Lab Diet 5LJ5 and water containing 16 mg/L NTBC (2-(2-nitro-4-fluoromethylbenzoyl)- 1,3-cyclohexanedione). 1 day prior to transplantation, 1.25 x10^9^ pfu/mL of *<u>u</u>rokinase type <u>P</u>lasminogen <u>A</u>ctivator* (*uPA*)-expressing adenovirus particles (Yecuris) were retroorbitally injected into mice and 0.5-1.5x10^6^ hPSC-derived day 18 hepatocytes were intrasplenically transplanted. 1 day before transplantation, liver injury was induced by cyclical withdrawal of NTBC for 5 to 7 days and then provision of 8 mg/L NTBC for 2 to 3 days.

### Surface marker FACS

Cells were dissociated using TrypLE Express, pelleted, resuspended in FACS buffer (0.5% BSA Fraction V + 5mM EDTA in 1 x PBS) and strained through a 40μm filter (BD Biosciences) to generate a single-cell suspension. The cell suspension was aliquoted into individual tubes and stained with α-CD99 PE, α-CD99-APC, α-CD10 APC, α-CD201-APC or α-CXCR4-PE-Cy7 (**Table S2**) for 30 minutes on ice in the dark. Subsequently, cells were washed 3 times with 1-2 mL of FACS buffer, resuspended in 300 μL of FACS buffer containing 100 ng/mL DAPI into a FACS tube and analysed by FACS using BD LSR Fortessa X20.

## DATA AVAILABILITY

Microarray data are available from the NCBI GEO repository (accession GSE98324). Data will be made publicly available upon acceptance.

## AUTHOR CONTRIBUTIONS

L.T.A. designed the experiments. L.T.A., A.K.Y.T., S.H.G., S.H.C. and J.J.H.L. differentiated hPSCs, and with Q.F.C., transplanted hPSC-derived hepatocytes. A.M.I. performed Western blotting and generated the reporter line with B.F.P. K.L.L. generated and analyzed microarray data. K.M.L., A.C., I.L.W. and L.T.A. conducted surface marker analyses. L.T.A., K.M.L. and B.L. oversaw the study and wrote the manuscript.

## ACKNOWLEDGEMENTS

We dedicate this paper to the memory of Professor Lorenz Poellinger and thank him for his generosity and support. We are grateful to Massimo Nichane, Wai Leong Tam, Vince Luca, Dahai Zheng, Edwin Lee, Danial Asraf, Farid Juraimi, Roger Foo, Fabio Michelet, Wencai Zhang, Malin Boezelman, Hui Cheng, Sivakamasundari Vijayakumar, May Yin Lee, Keefe Chee, Benjamin Chua and Shyh Chang Ng for kindly sharing reagents and support. We thank Rani d/o Ettikan for logistical support and thank Bruce Wang, Wen Chuan Peng, Philip Beachy, Chiea Chuen Khor and Huck Hui Ng for discussions. This study was supported by the Agency for Science, Technology and Research (A*STAR) through ETPL Gap Funding Grant ETPL/14-R15- 009 to L.T.A. and through the Siebel Investigator Program to K.M.L.

## SUPPLEMENTARY INFORMATION

### Cell culture

Human ESC lines H1, H7, H9, HES2, HES3, ESI035, H1 *FAH-Clover* knock-in reporter line, H9 *EF1A-BCL2-2A-GFP* and H9 *EF1A-BCL2-2A-IRES*-*UBC-tdTomato-2A-Luciferase* (Loh et al., 2016) and human iPSC line BJC1 were maintained in feeder-free conditions using mTeSR1 medium (StemCell Technologies, 05851/2). Cell lines that are confirmed to be mycoplasma-free before differentiation. Anterior primitive streak was specified using Definitive Endoderm Induction Medium A for 24 hours, followed by Definitive Endoderm Induction medium B for 24 hours to induce definitive endoderm (Gibco, A27654SA). Midgut or hindgut cells were specified from DE using 50 to 100 ng/mL FGF2, 10 ng/mL BMP4 and 3 μM CHIR99201 in base medium CDM2 for 4 days (Loh et al., 2014). Early pancreatic endoderm cells were specified from DE using 1 μM C59, 250 nM DM3189, PD0325901, 2μM ATRA in CDM3 for 1 day followed by 10 ng/mL ACTIVIN, 1 μM C59, 250nM DM3189, PD0325901, 2 μM ATRA and 150 nM SANT1 in CDM3 for 3 days. The composition of CDM2 base medium has been previously reported (Loh et al., 2014; Loh et al., 2016; Touboul et al., 2010). CDM3 base medium comprises 10% KOSR, 0.1% PVA, IMDM/F12 (1:1), 1% concentrated lipids and 1% Pen Strep. CDM4 base medium comprises of amino acid supplements, 15 μg/mL Transferrin, 1% Glutamax, IMDM/F12 (1:1), 1% Pen Strep and 1% concentrated lipids. Amino acids were purchased from Sigma Aldrich, dissolved as per manufacturer’s recommendation, and combined to form amino acid-rich concentrated 10x medium. These amino acid supplements include glycine, L-alanine, L-arginine hydrochloride, L-asparagine, L-aspartic acid, L-cysteine hydrochloride, L-histidine hydrochloride, L-Lysine hydrochloride, L-methione, L-phenylalanine, L-proline, L-serine, L-threonine and L-valine. Finally, CDM5 base medium is similar to the CDM4 base medium except that it does not contain additional amino acid rich mixture. To generate day 12 hepatic progenitors, hPSC-derived liver bud progenitors were treated with Dexamethasone (10 μM), RO4929097 (2 μM) or DAPT (10μM), forskolin (10 μM), ± human recombinant insulin (10 μg/mL), ± BMP4 (10 ng/mL), ± Oncostatin M (10 ng/mL) and ± ascorbic acid-2 phosphate (AAP; 200 μg/mL) for 6 days. To generate day 18 hepatocytes, hPSC-derived hepatic progenitors were treated with Dexamethasone (10 μM), RO4929097 (2 μM) or DAPT (10 μM), forskolin (10 μM), human recombinant insulin (10 μg/mL), and ascorbic acid-2 phosphate (AAP; 200 μg/mL). Cryopreserved human hepatocytes (Thermofisher Scientific, HU1352, HU8055 or HU8132) were thawed with CHRM (Gibco, CM7000), plated and grown in matrigel sandwich cultures (Gibco, CM3000, CM4000) as per manufacturer’s instructions.

### Design and construction of Cas9 plasmids

Primers used for constructing the plasmids are listed in **Table S2**. All restriction enzymes were purchased from NEB, unless stated otherwise. PCR reactions were conducted using Q5® Hot Start High-Fidelity 2X Master Mix (NEB, M0494L) or Phusion High-Fidelity PCR Master Mix with HF Buffer (ThermoFisher Scientific, F531S). Ligations were conducted using isothermal assembly with NEBuilder® HiFi DNA Assembly Master Mix (NEB, E2621L) or In-Fusion® HD EcoDry™ Cloning Plus (Clontech, 638915). Primers and dsDNA fragments were ordered from Integrated DNA Technologies (IDT). EF1α promoter was PCR amplified from N205 plasmid (Addgene plasmid # 44017), a gift from Jerry Crabtree (Hathaway et al. 2012). Amp pUC fragment was obtained from pCMV-Bsd (Thermo, V51020). Plasmid pX330 (Addgene plasmid # 42230), a gift from Feng Zhang (Cong et al. 2013), was digested with XbaI & AarI (Thermo, ER1582) and ligated with EF1α promoter. Next, the modified plasmid was digested with SspI and XbaI and then ligated with the Amp pUC fragment. The BmsBI chimaeric gRNA cassette was amplified from a gBlock based on px335 (Addgene plasmid # 42335), a gift from Feng Zhang (Cong et al., 2013). The cassette was subsequently ligated into XbaI cut site of our modified Cas9 plasmid. To add a 2A-linked mRuby2, the plasmid was first digested with PmlI and EcoRI, and the Cas9 3’ fragment was amplified from px330 and 2AmRuby2 (Lam et al., 2012) fragment was amplified from a gBlock. Next, these two PCR fragments were ligated with the digested plasmid. Enhanced specificity of Cas9 was attained by specific mutations of the Cas9-protein sequence (Kleinstiver et al., 2016; Slaymaker et al., 2015). Such mutations (K848A/K1003A/R1060A) were introduced to our Cas9 plasmid by cloning in a gBlock. 5’ fragment of Cas9 and 3’ fragment plus 2AmRuby2 were amplified from the plasmid, and mutated sites from gBlock. The plasmid was digested with FseI and EcoRI and fragments ligated in the digest. hPSCs are very sensitive to DNA damage (Momcilovic et al., 2009; Momcilovic et al., 2010) and we found that Cas9 targeted hPSCs had low survival and low number of correctly targeted clones. Inhibiting the TP53 checkpoint could increase survival of targeted hPSCs (Ihry et al., 2017). mtp53 dominant negative fragment was amplified from gBlock based on pCE-mp53DD (Addgene plasmid # 41856), a gift from Shinya Yamanaka (Okita et al. 2013) and ligated it into our Cas9 plasmid digested with EcoRI to generate our final construct pMIA3 1sg-eSpCas9-2AmRuby2-2Amp53DD”.

### Design of gRNA

The genomic sequence of the end of human FAH CDS (chr15:80186000-80187000) was uploaded to Benchling (https://benchling.com/) and single gRNAs were designed using the online search algorithm. One gRNA (GAGCAGAGAAAATCTCATGA, negative strand) that overlaps the *FAH*-stop codon was selected. Oligos with the gRNA sequence and the complementary sequence (TCATGAGATTTTCTCTGCTC), with CACC- and AAAC- added to the 5’ end of each oligo, were purchased from IDT. Finally, the oligos were annealed and ligated into BsmBI-digested pMIA3 1sg-eSpCas9-2AmRuby2-2Amp53DD.

### GFP reconstitution assay

The gRNA cutting efficiency was confirmed by GFP reconstitution assay using pCAG-EGxxFP plasmid (Addgene plasmid # 50716), a gift from Masahito Ikawa (Mashiko et al., 2013), as described previously. Briefly, the target sequence was amplified from H1 hPSCs genomic DNA and then cloned into the SalI cut site on pCAG-eGxxFP (for primers see **Table S2**). The plasmid was then transfected into HEK293T cells with or without FAH-pMIA3. 48h later, strong GFP signal was observed when both plasmids were transfected indicating Cas9 cleavage activity (**Fig. S5a**,**b**,**c**).

### Design and construction of donor plasmids

Our homology directed repair donor plasmid was derived from OCT4-2A-eGFP-PGK-Puro (Addgene plasmid # 31938), a gift from Rudolf Jaenisch (Hockemeyer et al., 2011). 3’ homology arm for human *FAH* was amplified from gDNA from H1 hPSCs. The amplified product was then ligated into OCT4-2A-eGFP-PGK-Puro plasmid that had been double digested with AscI & FseI. To swap the resistance gene from puromycin to blasticidin, the mPgk1 promoter & polyA fragments were first amplified from OCT4-2A-eGFP-PGK-Puro and Blasticidin (Bsd) resistance gene from pCMV-Bsd plasmid. These fragments were ligated into BsrGI & AscI double-digested donor plasmid to generate a construct containing OCT4-2A-eGFP-PGK-Bsd-FAH. The modified plasmid was then digested with NheI & PacI to allow ligation of P2A-linked Clover fluorophore (Lam et al., 2012), that had been amplified from a gBlock. The PGK promoter was amplified from OCT4-2A-eGFP-PGK-Puro, to make OCT4-2A-Clover-PGK-Bsd-FAH. A Gly-Ser-Gly sequence was added to the start of the P2A sequence as this has been shown to increase the 2A-peptide cleavage efficiency (Kim et al., 2011; Szymczak et al., 2004). The Bsd CDS was amplified from pCMV-Bsd and the negative selection agent thymidine kinase (TK) CDS from pLOX-TERT-iresTK (Addgene plasmid # 12245), a gift from Didier Trono (Salmon et al. 2000). The two PRC fragments were then ligated into PacI & AflIII digested donor plasmid, to generate the construct OCT4-2AClover-PGK-Bsd2ATK-FAH. NotI was then used to digest this plasmid and a PGK-DTA negative selection cassette was ligated outside the homology arms, amplified from OCT4-2A-eGFP-PGK-Puro and a gBlock. Finally the 5’ homology arm of human *FAH* was amplified from H1 hPSC gDNA and then ligated into the SbfI- & NheI-digested plasmid to complete the FAH-2AClover-PGK-Bsd2ATK-FAH-PGK-DTA donor plasmid.

### Generation of FAH-2AClover H1 hES reporter cell line

1.5x10^6^ H1 hPSCs were used in one reaction of an Amaxa nucleofection (Lonza) targeting. 5 g of both FAH-pMIA3 and FAH-2AClover-PGK-Bsd2ATK-FAH-PGK-DTA donor plasmid were nucleofected with the P3 Primary Cell kit (V4XP-3024) using CM-113 program. Prior to nucleofection the cells were pretreated with ROCK-inhibitor (Y-27632, Tocris) for at least 1 hour and then dissociated into a single-cell suspension with StemPro Accutase (A1110501, ThermoFisher). After nucleofection, the cells were seeded in mTeSR supplemented by CloneR (05888, Stem Cell Technologies). After 24h, the media was changed to mTeSR and cells were left to recover. At approximately 48-72h, Bsd (A1113903, ThermoFisher) was added to the media at 10μg/ml to select for targeted cells. Individual colonies were manually picked from wells after 10-14 days of selection and further expanded for screening.

12 clones were picked from the targeted wells. 11 clones survived the expansion and gDNA was isolated for screening. Primers for amplifying the FAH target region in the eGxxFP plasmid (also annotated “eGxxFP primers” were used for screening the targeted clones alongside FAH- clover forward and reverse primers (**Table S2**). Expected PCR product size for wild-type cells is 422 bp with eGxxFP primers and for targeted cells 2.3 kbp with “FAH-clo” primers. 10 out of 11 clones picked amplified PCR bands at both WT and targeted sizes, suggesting they were heterozygous clones (**Fig. S5d**,**e**). 3 out of the 10 clones had no mutations on the WT allele as confirmed by Sanger sequencing (**Fig. S5g**).

Heterozygous clones that were confirmed were further expanded and targeted with 10μg of pCAG-Cre:GFP (Matsuda & Cepko 2007) (Addgene plasmid # 13776), a gift from Connie Cepko. These cells were allowed to recover for 48h after which they were sorted for GFP. GFP^+^ cells were plated in CloneR supplemented mTeSR at a density of 100 to 1000 cells per well in a 6- well tissue culture plate. After 48h, the media was changed to mTeSR and clones were allowed to expand for 10-14 days. Clones were then manually picked for expansion and screening. PCR screening and Sanger sequencing was done with eGxxFP primers as above. Expected PCR amplicon size for correctly targeted allele is 1.3 kbp (**Fig. S5f**). Next, confirmed clones were expanded and used in downstream experiments.

### Intracellular FACS

AFP antibodies (DakoCytomation, A000829) were conjugated with R-phycoerythin (rPE) using rPE labeling kit (abcam, ab102918). Cells (either undifferentiated hPSCs or hPSC-derived liver cells) were dissociated into single cells using TrypLE Express (Gibco, 1260413) and centrifuged at 2000 rpm for 3 minutes. Each cell pellet was washed with 1x PBS (Gibco, 14190235), strained using a 100μm strainer (Falcon, 08-771-19) and counted using a hemocytometer before fixation in BD Cytofix/Cytoperm buffer (BD Bioscience, 554714) on ice for 20 minutes. Next, fixed cells were washed 2 times with 2 mL 1x BD Perm/Wash buffer (pre-warmed to room temperature) (BD Bioscience, 554723) at room temperature and pelleted. The cell pellet was then resuspended in 1x BD Perm/Wash buffer and 100μLof cell suspension was aliquoted into individual tubes for separate stains. Stained or unstained controls were included. Either anti-ALB-APC (R&D, IC1455A, 0.4μL per 150,000 cells) or anti-AFP-PE (0.33μL per 150,000 cells) was used. Anti-AFP-PE antibody/cell mixture was incubated at room temperature for 30 minutes in the dark while anti-ALB-APC antibody/cell mixture was incubated at room temperature for 20 minutes in the dark. Subsequently, the unstained or stained cells were washed 2 times with 2 mL 1x BD perm/wash buffer at room temperature and pelleted. The pellet was then resuspended in 300μL 1x BD perm/wash. FACS run was performed using BD LSR Fortessa X20 and FACS data were analyzed using FlowJo.

### Sorting of Clover^+^ and Clover^-^ cells by FACS

Cells were dissociated as single cells and stained with DAPI prior to FACS. Separate samples of FAH-Clover^+^ and Clover^-^ populations were gated and collected and then harvested for gene expression analyses.

### High-throughput antibody screens

BD Lyoplate Screening panel (BD Biosciences, 560747) antibodies (**Table S1**) were reconstituted with deionized water before their use to stain cells following manufacturer’s instructions. Cell suspension was filtered through 100 μm strainer to remove clumps. 75 μL cell suspension was added to each antibody-containing well, pipette-mixed 3 times and incubated in the dark at 4°C for 20-30mins. The cells were pelleted at 500g for 6 minutes and supernatant was discarded by plate inversion. The cells were then washed twice with 200 μL cell staining buffer by pipette mixing, resuspended in DAPI-containing cell staining buffer and analysed with BD LSR Fortessa X20.

### Immunostaining and image analyses

Cells were washed once gently with 1x PBS, fixed with 4% formaldehyde in PBS for 15-20 minutes at room temperature, washed 3 times with 1x PBS and blocked with blocking buffer (10% Donkey Serum + 0.1% Triton X in 1x PBS) for 1 hour at room temperature. Next, the cells were stained with antibodies diluted in 1% Donkey Serum + 0.1% Triton X in 1x PBS at 4°C overnight (see **Table S3** for all antibodies used). The next day, the cells were washed 3 times with washing buffer (0.1% Triton X in 1x PBS) and then stained with fluorophore-conjugated secondary antibodies diluted in at 1:1000 (1% Donkey Serum + 0.1% Triton X in 1x PBS) in the dark at room temperature for 1 hour. Finally, the cells were washed once with 100 ng/mL DAPI/PBS once and twice with 1x PBS before visualization using Zeiss Axio Vision. “No primary antibody staining” or undifferentiated hPSC negative controls were used to adjust exposure times for minimal background fluorescence detection. The stained cells were counted using Image J (Abràmoff and Magalhães, 2004; Collins, 2007). Image preprocessing including gray scale conversion, threshold setting, image segmentation and noise reduction were performed.

### Western blotting (WB)

Cell samples were lysed using radioimmunoprecipitation assay (RIPA) buffer supplemented with protease inhibitor cocktail (Nacalai Tesque, 25955-11). Protein concentration was determined using Pierce BCA Protein Assay (ThermoScientific, 23225). 40 μg of protein lysates were separated by SDS-PAGE and transferred onto a PVDF membrane (100V at 4°C, for approximately 1 hour). Membranes were blocked with 5% milk in TBS for 1 hour at room temperature. Membranes were briefly washed with Tris-buffered saline (TBS) and after this incubated with the primary antibody (see **Table S3** for all antibodies used) at 4°C overnight. The following day, membranes were washed 3 times with TBS and incubated with the appropriate HRP-conjugated IgG secondary antibody for 2 hours at room temperature. All antibodies, primary and secondary, were used with 5% milk in TBS. Following incubation with secondary antibody, the membranes were washed 3 times in TBS. Washed membranes were developed using SuperSignal West Pico ECL (Thermo Scientific, 34077) and imaged with BioRad ChemiDoc MP Imager.

### Immunohistochemistry (paraffin)

Immunohistochemistry was performed on paraffinized murine tissue that was embedded onto glass microscope slides and sectioned using a microtome. Sections were deparaffinized through a series of 5-minute washes with xylene, 100% ethanol, 95% ethanol, 70% ethanol and lastly milli-Q water. Endogenous peroxidase activity in sections were blocked using blocking buffer (65 mL of 100% methanol, 3.5 mL of 30% hydrogen peroxidase (Sigma) and 31.5 mL of milli-Q water) for 30-minute at room temperature. Finally, antigen retrieval was performed at boiling temperature for 30 minutes in a 10% pH 6-citrate buffer.

The tissues were subsequently blocked with donkey serum for 1 hour at room temperature in a hydration chamber. Sections were stained with primary antibodies (Vector Laboratories, Vectastain ABC kit; PK-6101) at 4°C overnight, later washed 3 times with 0.1% Triton in 1x PBS done in 5-minute incubation intervals (with fresh buffer applied during each wash) and stained with secondary biotinylated antibody at room temperature for 30 minutes. Thereafter, the samples were lightly rinsed with 1x PBS for 5 minutes on an orbital shaker. 250 μl of ABC reagent was added to each sample and left to incubate at room temperature for 1 hour. The slides were then washed 3 times with 1x PBS for 5 minutes.

For peroxidase detection, DAB kit (Vector Laboratory, SK-4100) was used for substrate binding. The substrate concoction (5 mL of milli-Q water, 2 drops of buffer stock, 4 drops of DAB and 2 drops of hydrogen peroxide) was applied onto the samples and left to incubate for 5 minutes at room temperature. After the 5 minutes incubation, the samples were rinsed 3 times with milli-Q water, changing fresh water each time. The samples were then placed into a vertical plastic chamber containing hematoxylin and incubated for 20 minutes.

After the hematoxylin counterstaining step, the samples were rinsed 3 times with milli-Q water before proceeding to the dehydration steps. Dehydration steps comprises: 70% ethanol, 95% ethanol, 100% ethanol and xylene wash steps, all with 5 minutes incubation. The counterstained samples were permanently mounted using Vectamount reagent (H-5000) from Vector Laboratory.

### Immunohistochemistry (cryoblock)

Mouse liver tissue was fixed in 4% formaldehyde diluted in 1x PBS at 4°C overnight and then immersed in 30% sucrose solution for cryoprotection. The liver tissue was then embedded in Optimal Cutting Temperature (OCT), sectioned and deposited onto microscope slides (A*STAR IMCB histology lab). The microscope slides were thawed in room temperature and stained following the immunostaining method described above.

### Gene expression profiling by RT-qPCR

RNA was extracted using Zymo RNA extraction kit. Reverse transcription was performed with random primers (Applied Biosystems RT kit) to generate cDNA. Gene expression was quantified using gene-specific primers and Quantitect SYBR Green master mix (Qiagen) or KAPA SYBR® Fast qPCR kit (KAPA Biosystems, KK4602). NCBI primer designing tool was used to design gene-specific primer sequences. Primers used were validated for their specificity and those with efficiency between 90-110% were used (**Table S4**). Gene expression profile was analysed using Microsoft excel and heatmaps were generated using GenePattern version 2.0 (Reich et al., 2006).

### Microarray

Total RNA was extracted from hPSCs and hepatic derivatives using the RNeasy Micro kit (Qiagen) and profiled for RNA integrity using the 2100 Bioanalyzer (Agilent). 500ng of high purity and high integrity samples with 260/280 and 260/230 absorbance ratios >1.9 and RNA integrity numbers (RIN) >8.0 were reverse transcribed into cDNA and in vitro transcribed into biotin-labelled cRNA using the TargetAmp-Nano labeling kit for Illumina Expression BeadChip (Epicentre). The cRNA was hybridised on HumanHT-12 v4 Expression BeadChips and scanned on the HiScan system (Illumina) according to the manufacturer’s specifications. The raw microarray data was background subtracted using the BeadStudio Data Analysis Software v3.1.3.0 (Illumina) and normalised using the cross-correlation method (Chua et al., 2006). Differential gene expression was defined based on a fold-change cutoff of >1.5 compared to the average of the undifferentiated hESC baseline controls (**Table S5**). Heatmaps of the fold-change in gene expression on a Log 2 scale were generated using Cluster and TreeView. Gene ontology analyses on the genes downregulated upon removal of DAPT, Forskolin, Ascorbic acid and Insulin were conducted using DAVID/EASE (Huang et al., 2008; Huang et al., 2009). Background subtraction using Illumina HT-12 v3 database was chosen in the settings.

### Animal husbandry and blood sampling

The IACUC and IRB committee had approved all procedures performed in the study. FRG mice on NOD or C57BL6 background (Yecuris, 10-0008 or 10-0001) were handled and housed under aseptic conditions. Each mouse had its ears notched for long-term tracking. The mice were fed ad libitum with irradiated LabDiet 5LJ5 chow formulated with high fat and low protein content to avoid excessive tyrosine levels, which can lead to liver damage. NTBC (Yecuris, 20-0026) was dissolved in sodium carbonate to generate 1 mg/mL stock solution. A final dose of 16 mg/L was given to breeders or non-experimental mice, whereas 0 to 8 mg/L NTBC was provided to experimental mice on selected days. All experimental mice were treated with the same NTBC cycling condition. 3% Dextrose was added to the drinking water to offset the bitter taste of NTBC. Antibiotics such as Sulfamethoxazole (SMX) and Trimethoprin (TMP) (Yecuris, 20-0037) were added to the water once every other week for 2-4 days to prevent infection in the immune-compromised mice. To aid in caloric intake, each cage was supplemented with a dish of liquid diet, prepared by adding 1 volume of STAT high caloric liquid (PRN Pharmaceutical, G8270) to 1 volume of 3% Dextrose (Sigma) drinking water. Masses of FRG mice were measured weekly. Blood was collected using lancets for submandibular bleeding and allowed to coagulate in 4°C overnight. The next day, serum was harvested by centrifugation and removal of red blood cells.

### Retro-orbital injections, intrasplenic and intrahepatic injections

Adenovirus particles expressing urokinase Plasminogen activator (uPA) (Yecuris, #20-0029) were injected at a dose of 1.25*10^9^ pfu at a volume of 100 μL per 25g mice approximately 24 hours before transplantation. Pertaining to intrasplenic or intrahepatic injections, 20-300 μL of cells was injected into the spleen or liver of 4-6 week-old FRG mice using 26G-31G needles.

### Intrahepatic injections into neonate and adult livers

For neonatal intrahepatic transplantations, 0-48h old FRG pups were injected with approximately 200,000 cells directly into the liver using 31G needles. The pups were then rubbed with bedding and returned to the cages. Pertaining to intrahepatic transplantation into adult livers, 50 μL of cells were injected into multiple sites within the liver using 26G-31G needles. Bleeding was stopped by gently applying pressure on the puncture site upon withdrawal of needle.

### Bioluminescence imaging

Mice were pre-transplanted with hPSC-derived hepatocyte-like cells, which overexpress *luciferase* gene. Prior to bioluminescence imaging, the transplanted mice were anesthetized with 1 to 3% isoflurane-mixed oxygen. To minimize imaging signal interference, mice were depilated at abdominal region to fully expose skin surface as hair will absorb and scatter light which may result in lower signal output. Mice were then weighed to determine their weight. D-luciferin solution was reconstituted by adding 30 mL of saline/1 x PBS to 1g of D-luciferin stock (Promega, InvivoGlo) and then filtered. 167 μg/g firefly D-luciferin was injected intraperitoneally into each mouse. After 10min of incubation time, the mice were positioned in the Perkin Elmer IVIS Spectrum facing upright (0% relative to supine position) in which the whole liver region was exposed to the camera’s field of view. The parameters of bioluminescence imaging were as follow: Exposure – 40s; binning – medium; excitation – block; emission – open; structure – no; FOV – D; Fstop – 1; and height – 1.5.

### Enzyme linked immunosorbent assay (ELISA)

ELISA Accessory kit (Bethyl Laboratories, Inc, E101) was used to quantify human serum albumin levels. Assay was performed as per manufacturer’ instructions. Absorbance was measured using an ELISA plate reader at 450 nm.

### Bilirubin quantification

Bilirubin levels in mouse serum were measured using a colorimetric Bilirubin assay kit (Sigma, MAK126). Total, direct or blank working reagents were prepared as per manufacturer’s instructions. 20 μL serum was added per well and total, direct or blank working reagents were added to each sample. Colorimetric product was measured at 530 nm using the Sunrise^TM^ microplate reader. Bilirubin concentration was then calculated using the following formula [(A530sample – A530blank) x 5 mg/dL]/ [A530 calibrator – A530 water].

### Survival curve analyses

Mice were checked for survival daily and Kaplan-Meier survival analysis was conducted using GraphPad Prism v7.00 for Mac (GraphPad Software, La Jolla, CA, USA). Statistical analyses were performed using the one-sided Mantel-Cox log-rank test. Data from 3 independent transplantation experiments were analyzed.

### Statistics

No statistical method was used to pre-determine sample size for *in vitro* or *in vivo* experiments. Experiments were not randomized. The investigators were not blinded to the allocation during experiments or outcome assessment.

**Figure S1.**
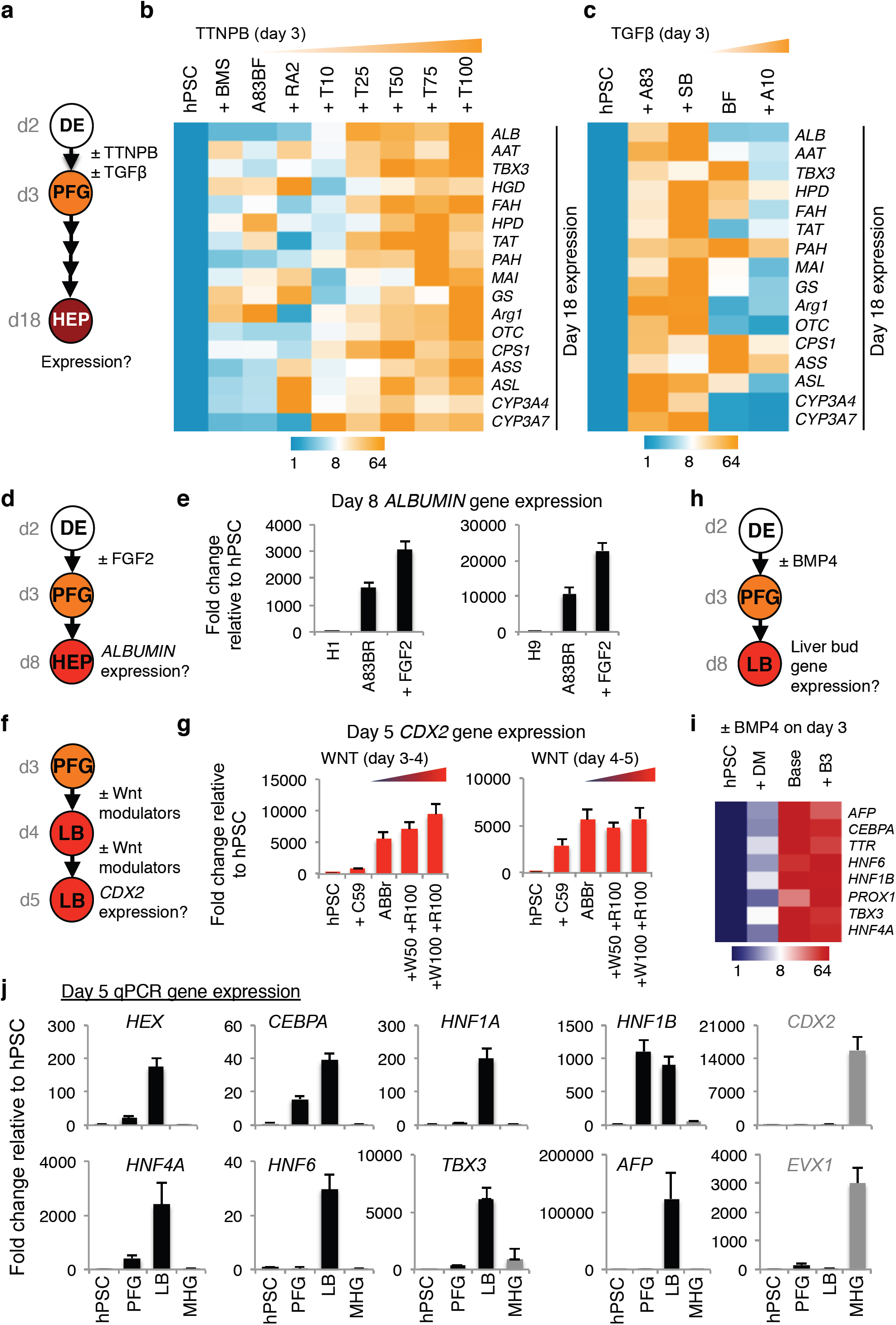
Regulation of early foregut competence. **a)** Experimental strategy to treat DE with RA or TGFβ modulators on the day 2-3 interval and evaluating its subsequent impact on day 18 hepatocyte gene expression as shown in subpanels **b**, **c**. **b**) qPCR gene expression of day 18 hepatocyte markers after inhibition (BMS) or activation of retinoid signaling (using ATRA, 2 μM or various doses of TTNPB, 10-100 nM) in the presence of base condition A83BF (A83BF: A8301, 1 μM; BMP4, 30 ng/mL; FGF2, 10 ng/mL) on the day 2-3 interval. **c**) qPCR gene expression of day 18 hepatocyte markers after inhibition (A8301, 1 μM or SB505124, 1 μM) or activation (ACTIVIN, 10 ng/mL) of TGFβ signaling in the presence of base condition BF (BF: BMP4, 30 ng/mL; FGF2 10, ng/mL) on the day 2-3 interval. **d**) Experimental strategy to treat definitive endoderm (DE) with FGF2 at 10 ng/mL) on the day 2-3 interval to produce day 3 posterior foregut (PFG) and assaying subsequent effects on hepatic gene expression by day 8, as shown in subpanel **e**. **e**) ALBUMIN qPCR of day 8 hepatic progenitors cells generated from endoderm treated on the day 2-3 interval with FGF2. **f**) Experimental strategy to treat posterior foregut (PFG) or liver bud progenitors (LB) with a WNT inhibitor (C59, 1 μM) or R-SPONDIN3 (R100, 100 ng/mL) and WNT3A at varying doses of 50 or 100 ng/mL on the day 3-4 or day 4-5 interval to produce day 4 or day 5 liver bud progenitors, as shown in subpanel **g**. **g**) CDX2 qPCR of day 5 liver bud progenitors cells generated from PFG treated on the day 3-4 interval with WNT modulators. **h)** Experimental strategy to treat DE with BMP4 or DM (DM3189) on the day 2-3 interval and evaluating its impact on day 6 liver bud differentiation, as shown in subpanel **i**. **i**) qPCR gene expression of day 5 liver bud progenitors after inhibition (DM) or activation of BMP signaling on day 2- 3 interval. **j**) qPCR gene expression of hPSC, day 3 hPSC-derived PFG and day 5 hPSC-derived liver bud (LB) and midgut/hindgut (MHG) progenitors.

**Figure S2.**
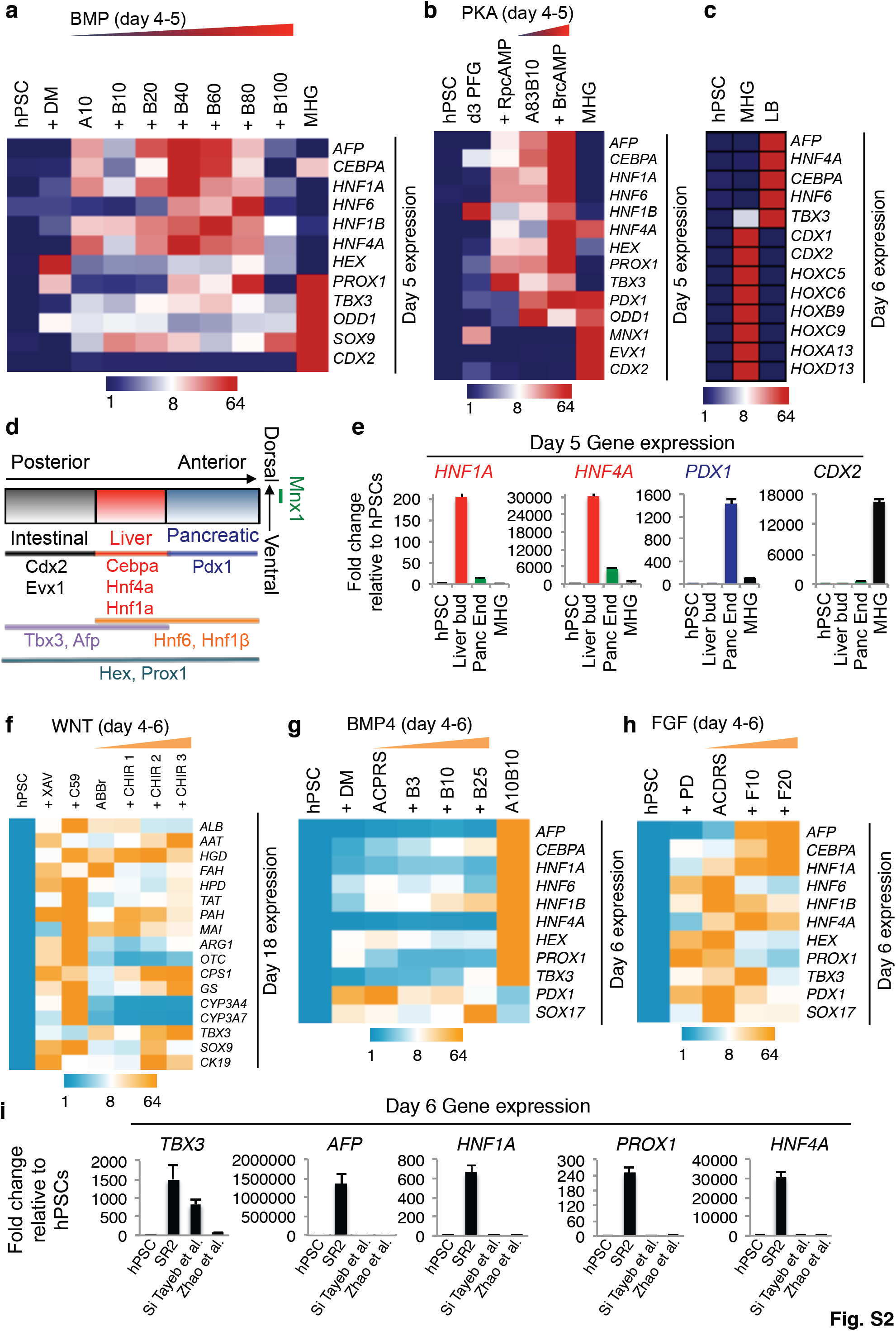
Lineage bifurcation between liver bud and pancreatic fates. **a)** qPCR gene expression of day 5 liver bud cells generated from endoderm treated on the day 4-5 interval with a BMP inhibitor (DM: DM3189, 250 nM) or varying doses of BMP4 (B, 10-100 ng/mL) in the presence of base condition A10 (A10: ACTIVIN at 10 ng/mL). MHG denotes hPSC-derived midgut/hindgut cells. **b**) Gene expression of day 6 liver bud cells after 2-day treatment of PKA inhibitor (rRpCAMP) or PKA agonist (BrCAMP) in the presence of base condition A83B10 (A83B10: A8301, 1 μM; BMP4, 10 ng/mL) during a day 4-5 interval and day 6 hPSC-derived MHG, as shown by qPCR. **c**) Gene expression of day 6 hPSC-derived liver bud and MHG cells derived as shown by qPCR. **d)** Cartoon of known organ domain markers expressed in ∼E8.5 mouse endoderm. **e**) Gene expression of day 5 liver bud progenitors, MHG or pancreatic endoderm cells (Panc End) derived from hPSCs as shown by qPCR. **f**) qPCR gene expression of day 18 hepatocytes generated from PFG cells treated on the day 4-6 interval with WNT inhibitors (C59, 1 μM or XAV939, 1 μM) or CHIR99201 (CHIR) of varying doses (1 μM, 2 μM or 3 μM). **g**) qPCR gene expression of day 6 cells after treatment of PFG cells on the day 4-6 interval a BMP inhibitor (DM: DM3189, 250 nM) or varying doses of BMP4 (3-25 ng/mL) in the presence of pancreatic inducing base condition (ACPRS = ACTIVIN + C59 + PD0325901 + ATRA + SANT1). A10B10: ACTIVIN, 10 ng/mL; BMP4, 10 ng/mL. **h**) qPCR gene expression of day 6 cells after treatment of PFG cells with a FGF/ERK inhibitor (PD032: PD0325901, 500 nM) or varying doses of FGF2 (10-20 ng/mL) in the presence of pancreatic inducing base condition (ACPRS = ACTIVIN + C59 + PD0325901 + ATRA + SANT1) on the day 4-6 interval. **i**) qPCR gene expression of H9 hPSC-derived liver bud progenitors generated from SR2 described in the present study and methods previously described in the literature (Si Tayeb et al., 2010; Zhao et al., 2012).

**Figure S3.**
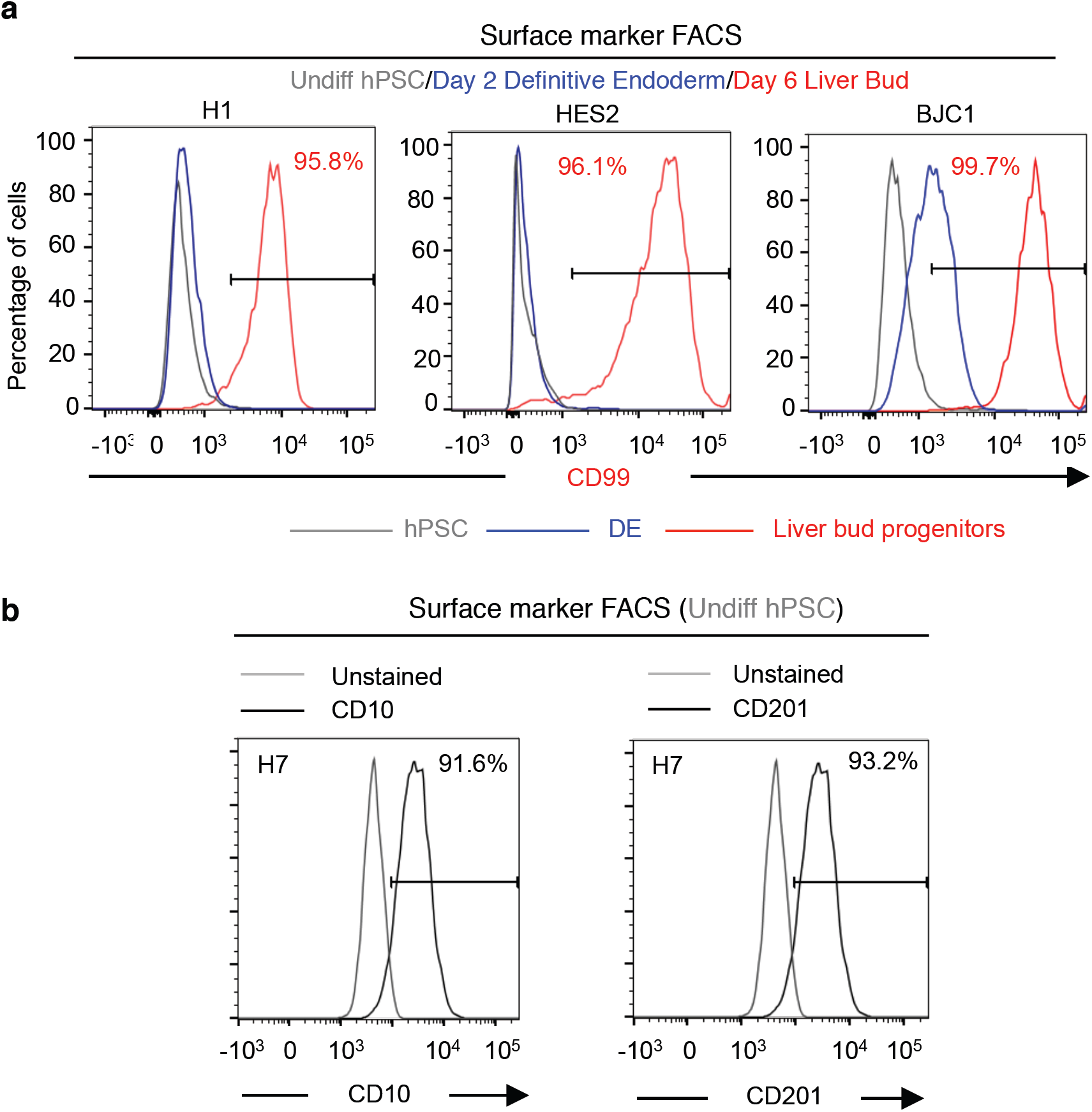
Identification of hPSC-, endoderm- and liver bud progenitor-specific cell surface markers using BD Lyoplate antibody screens. **a)** Percentages of H1, HES2 and BJC1 hPSC (grey), hPSC-derived endoderm (blue) and hPSC-derived liver bud progenitors (red) expressing surface marker CD99, as shown by FACS. (**b**) Percentages of H7 hPSCs expressing CD10 or CD201 as shown by live-cell FACS.

**Figure S4.**
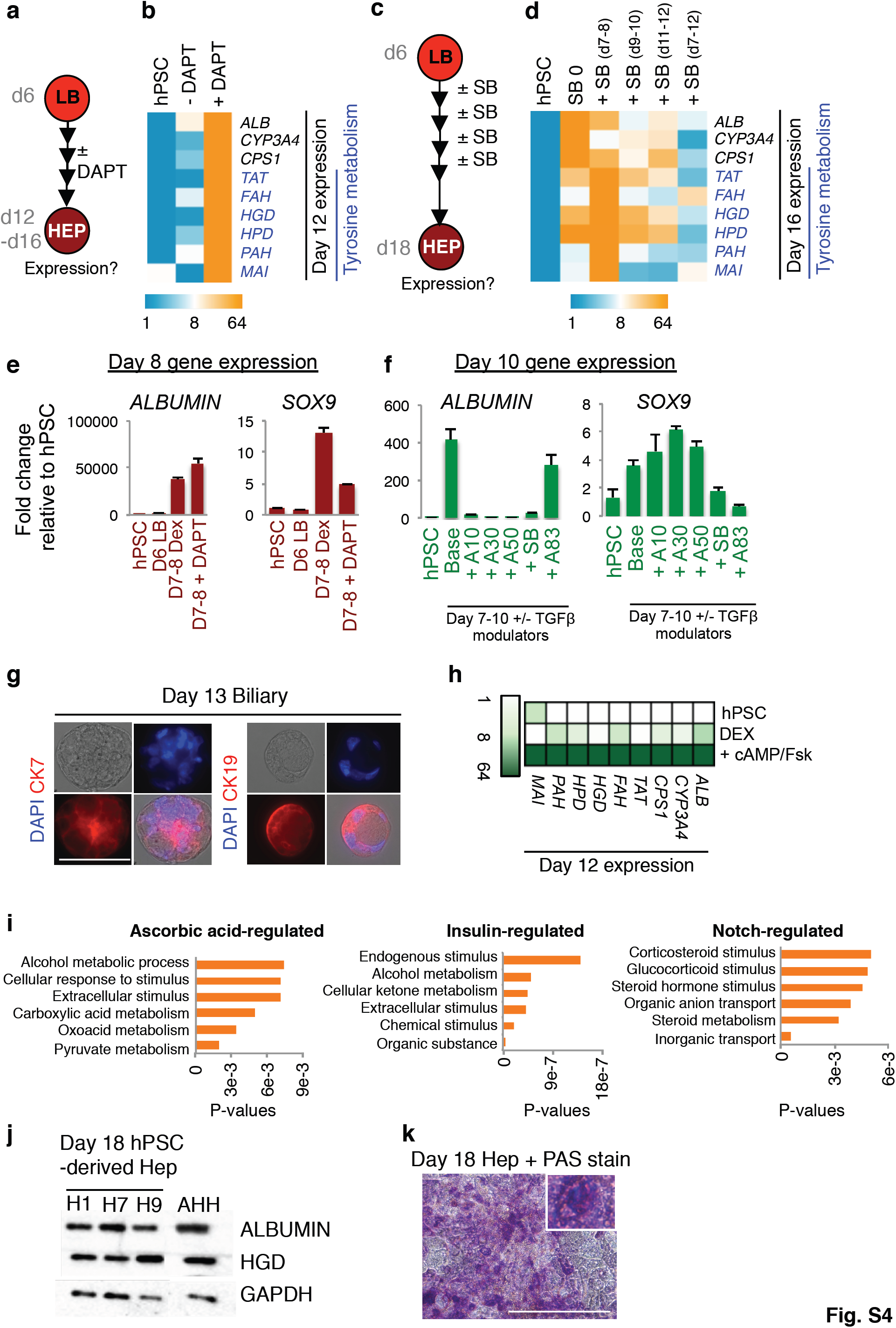
Signals inducing expression of tyrosine metabolic enzymes. **a)** Experimental strategy to treat day 6 liver bud progenitors (LB) with NOTCH modulators on the day 7-12 interval and assaying effects on hepatic gene expression by day 12, as shown in subpanel **b**. **b**) qPCR gene expression of day 16 hepatocytes generated from liver bud progenitors treated on the day 7-12 interval with a NOTCH inhibitor (DAPT, 10 μM). Each qPCR heatmap is representative of 4 independent experiments with technical duplicates. **c**) Experimental strategy to treat day 6 liver bud progenitors (LB) with a TGFβ inhibitor (SB505124, 1 μM) on the day 7-8, 9-10, 11-12 or 7-12 intervals and assaying downstream effects on hepatic gene expression by day 16, as shown in subpanel **d**. **d**) qPCR gene expression of day 16 hepatocytes generated from liver bud progenitors treated on the day 7-12, 12-16 or 7-16 intervals with a TGFβ inhibitor (SB505124, 1μM). Each qPCR heatmap is representative of 3 independent experiments with duplicates. **e**) qPCR gene expression of day 6 hPSC-derived liver bud (LB) progenitors before and after 2 day treatment of 10 μM DAPT in the absence or presence of 10 μM DEX. **f**) qPCR gene expression of day 10 hPSC-derived hepatic progenitors after 4-day treatment with TGFβ inhibitors A83 (A8301, 1 μM) or SB (SB505124, 1 μM) or TGFβ activator ACTIVIN at varying doses (10, 30, and 50 ng/mL) in the presence of base media. **g**) Immunostaining of CK7 and CK19 expression in day 13 hPSC-derived biliary progenitors, with DAPI nuclear counterstain DAPI, scale = 100μm. **h**) qPCR gene expression of day 12 liver cells after treatment with 8-BromocAMP (cAMP) and Forskolin (Fsk) in the presence of Dexamethasone (Dex). **i**) Gene ontology terms enriched amongst the genes regulated by DAPT, AAP or INS during the liver bud →→ hepatic progenitor differentiation step. **j**) Protein expression of ALBUMIN, HGD and GAPDH in hPSCs (undifferentiated H1, H7, H9), day 18 H1-, H7- and H9-derived hepatocytes and adult human hepatocytes (AHH) as shown by western blot. **k**) Day 18 hPSC-derived hepatocytes stained with Periodic Acid-Schiffs (PAS) stain; scale = 400μm. Each western blot is a representative of 2 independent experiments. Roadmap for accelerated human liver progenitor formation (Ang et al.), Pg. 40

**Figure S5.**
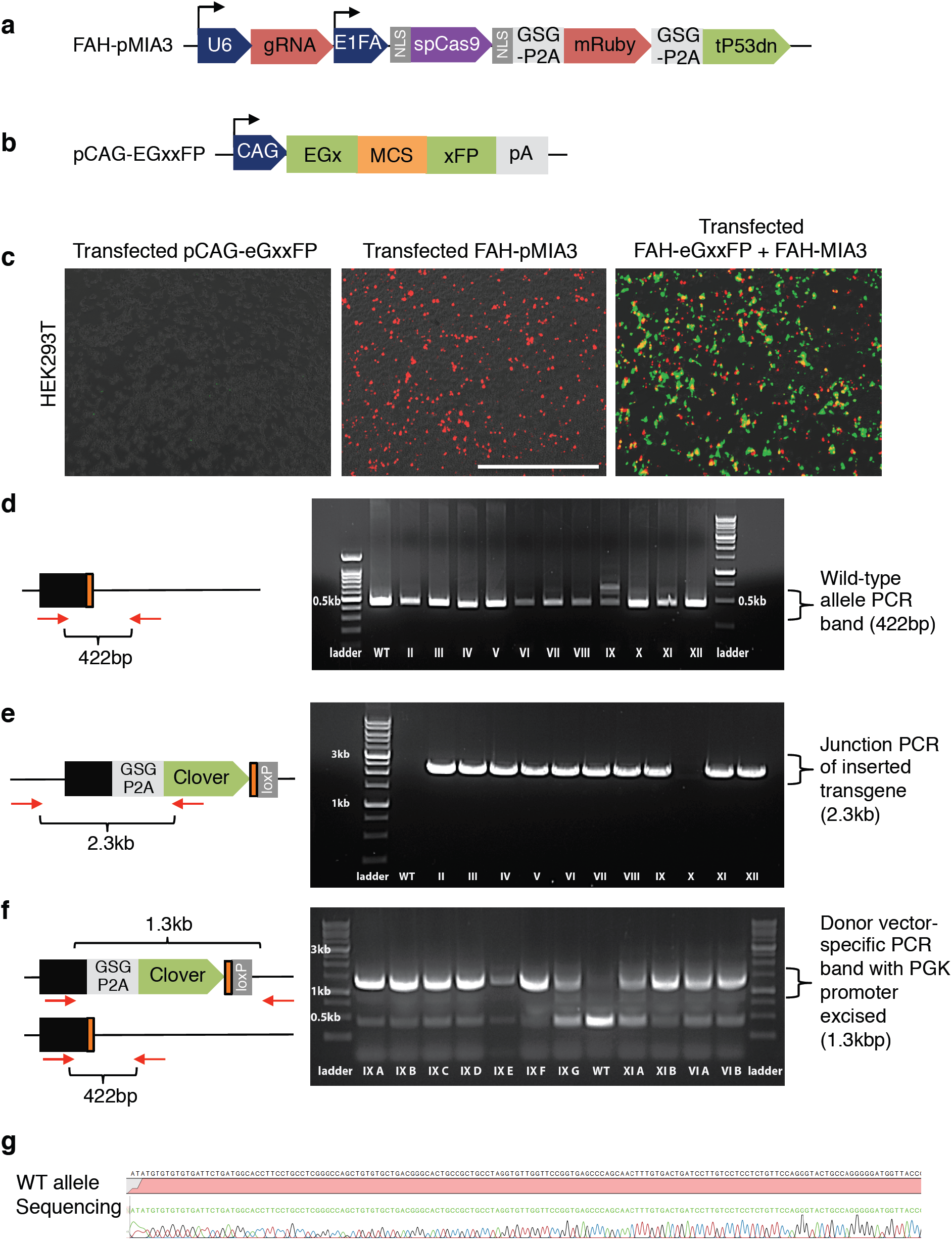
Generation of FAH-Clover hPSC reporter line by CRISPR/Cas9 mediated gene editing. **a**) Schematic of FAH-pMIA3 plasmid. **b**) Schematic of eGxxFP plasmid. **c**) Guide RNA and eSpCas9 cleavage efficacy test. HEK293T cells transfected with pCAG-eGxxFP plasmid, FAH-pMIA3 plasmid and both plasmids; scale = 1000μm. **d**) PCR screening of H1 FAH-2AClover clones with eGxxFP primers after Cre-excision. **e**) PCR screening of H1 FAH-2AClover clones with FAH-Clo junction primers. Arrows illustrates DNA regions that PCR screening primers bind to. **f)** PCR screening of H1 FAH-2AClover clones with eGxxFP primers. **g**) Sequencing of PCR products from WT allele of clones with no targeting on second allele with donor vector-specific primers.

**Figure S6.**
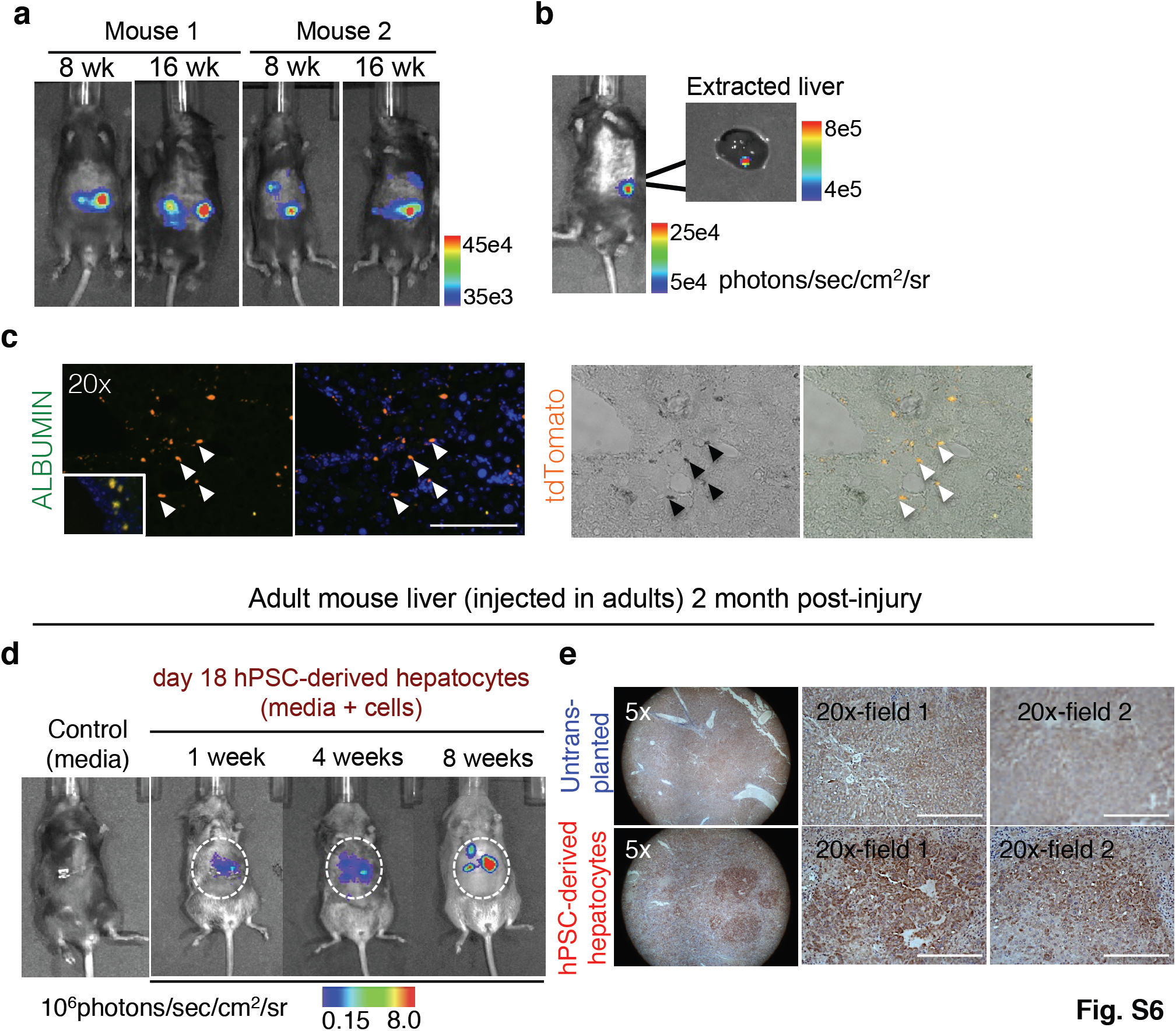
Tracking hPSC-derived liver cells in neonatal and adult *FRG* mice. **a)** *In vivo* tracking of 2 FRG mice with injected as neonates with hPSC-derived liver cells; bioluminescence imaging of the FRG mice 8 or 16 weeks (8 wks, 16 wks) after induced liver injury. **b**) Mouse with detected bioluminescence before liver extraction (left) and the extracted mouse liver (right); scale bar of signal intensity, in p/sec/cm^2^/sr. **c**) Mouse liver sections stained for human ALBUMIN (green) as shown by immunostaining; tdTomato-expressing cells appear red (arrowheads); scale bar =150mm. **d**) Bioluminescence imaging of mice injected with media as negative control, adult primary human hepatocytes and Luciferase+ hESC-derived liver cells (1, 4 and 8 weeks after injections). **e**) Untransplanted mice or mice transplanted with Day 18 hPSC-derived hepatocytes stained for ALBUMIN (brown) using human-specific ALBUMIN antibody at various magnifications as shown by immunohistochemistry; scale bar = 200μm.

**Table S1:** Surface marker expression on hPSC, hPSC-derived endoderm and hPSC- derived liver bud progenitors.

| Description | H7 UD | H7 D2 DE | H7 D6 LB | Description | H7 UD | H7 D2 DE | H7 D6 LB |
| --- | --- | --- | --- | --- | --- | --- | --- |
| abTCR | 0.22 | 0.00882 | 0.57 | CD314(NKG2D) | 11 | 0.00901 | 0.69 |
| B2-uGlob | 97.3 | 62.9 | 20 | CD32 | 0.064 | 0.00897 | 0.32 |
| BLTR-1 | 0.011 | 0 | 0.64 | CD321(F11 Rcptr) | 99.9 | 99.8 | 99 |
| CD10 | 87.6 | 64.3 | 2.84 | CD326 | 99.8 | 99.6 | 83.5 |
| CD100 | 60.2 | 0.38 | 0.71 | CD33 | 0.011 | 0.045 | 0.22 |
| CD102 | 19.2 | 0.91 | 1.28 | CD335(NKP46) | 0.074 | 0.027 | 1.48 |
| CD103 | 0 | 0.00957 | 0.23 | CD336 | 0.011 | 0.037 | 0.74 |
| CD104 | 6.72 | 0.00898 | 4.8 | CD337 | 0.096 | 0.00894 | 0.69 |
| CD105 | 17.9 | 5.09 | 0.66 | CD338(ABCG2) | 21.6 | 2.77 | 5.5 |
| CD106 | 0 | 0.00955 | 58.5 | CD34 | 0.011 | 0.019 | 30.2 |
| CD107a | 4.85 | 48.2 | 16.4 | CD340(Her2) | 100 | 100 | 99.9 |
| CD107b | 0.1 | 0.37 | 1.73 | CD35 | 1.75 | 0.13 | 0.35 |
| CD108 | 91.1 | 80.6 | 45.5 | CD36 | 0.043 | 0.045 | 0.78 |
| CD109 | 20.1 | 0 | 0.082 | CD37 | 0.011 | 0.027 | 0.24 |
| CD112 | 3.01 | 95.9 | 1.9 | CD38 | 0.31 | 0.063 | 0.55 |
| CD114 | 0 | 0.02 | 0.3 | CD39 | 0.032 | 0 | 0.42 |
| CD116 | 0.021 | 0.00999 | 0.36 | CD4 | 16 | 0.00959 | 0.13 |
| CD117 | 0.032 | 96.6 | 6.5 | CD40 | 60.5 | 57.5 | 1.75 |
| CD118 (LIF R) | 0.096 | 3.59 | 24.5 | CD41a | 0 | 0.00887 | 0.19 |
| CD119 | 99.9 | 68.2 | 82.9 | CD41b | 0 | 0.045 | 0.19 |
| CD11a | 0 | 0.011 | 0.27 | CD42a | 0.011 | 0.02 | 0.14 |
| CD11b | 0.011 | 0.00925 | 0.18 | CD42b | 0 | 0.027 | 0.29 |
| CD11c | 0.032 | 0.028 | 0.42 | CD43 | 43 | 0.13 | 0.92 |
| CD120a | 5.81 | 2.49 | 0.85 | CD44 | 95.8 | 48.8 | 13 |
| CD120b | 4.64 | 0.036 | 0.25 | CD45 | 0 | 0.00893 | 0.19 |
| CD121a | 0.011 | 0.00998 | 0.33 | CD45RA | 0.064 | 0.018 | 0.2 |
| CD121b | 0.022 | 0.04 | 0.54 | CD45RB | 0.05 | 0.018 | 0.34 |
| CD122 | 0.011 | 0.00992 | 0.23 | CD45RO | 8.77 | 0 | 0.074 |
| CD123 | 0.043 | 0.00971 | 0.25 | CD46 | 99.9 | 100 | 99.9 |
| CD124 | 0.15 | 0.00977 | 4 | CD47 | 100 | 99.6 | 98.4 |
| CD126 | 0.17 | 0.00976 | 2.52 | CD48 | 0.014 | 0.036 | 0.18 |
| CD127 | 0.032 | 0.04 | 0.44 | CD49a | 32.8 | 0.22 | 27.2 |
| CD128b | 2.01 | 0.02 | 0.32 | CD49b | 100 | 76.1 | 94.5 |
| CD13 | 4.37 | 23.1 | 33 | CD49c | 99.9 | 66.1 | 99.7 |
| CD130 | 2.48 | 36 | 60.4 | CD49d | 40.2 | 42 | 0.55 |
| CD132 | 0.011 | 0.054 | 0.14 | CD49e | 100 | 99.9 | 99.5 |
| CD134 | 0 | 0.02 | 0.79 | CD49f | 99.8 | 44.3 | 97.9 |
| CD135 | 0.043 | 0.049 | 0.3 | CD4V4 | 11.3 | 0.27 | 0.28 |
| CD137 | 0.011 | 0.03 | 5.49 | CD5 | 0.032 | 0.00959 | 0.3 |
| CD138 | 2.2 | 0.03 | 0.28 | CD50 | 96.8 | 38.8 | 0.96 |
| CD138 Ligand | 0 | 0.02 | 0.31 | CD51/61 | 0.036 | 10.9 | 4.69 |
| CD14 | 0.032 | 0.009 | 0.23 | CD53 | 0.036 | 0 | 0.27 |
| CD140a | 0.022 | 36.3 | 37.8 | CD54 | 90.6 | 68.9 | 54.1 |
| CD140b | 62.5 | 46.9 | 1.87 | CD55 | 100 | 100 | 97.1 |
| CD141 | 3.27 | 6.41 | 34.3 | CD56 | 99.9 | 83.2 | 55.9 |
| CD142 | 98.9 | 94.5 | 98.4 | CD57 | 99.4 | 99.7 | 93 |
| CD144 | 1.4 | 4.34 | 0.52 | CD58 | 100 | 100 | 99.6 |
| CD146 | 99.8 | 86.5 | 80.4 | CD59 | 99.9 | 100 | 99.9 |
| CD147 | 99.4 | 68.6 | 61.9 | CD6 | 0 | 0 | 0.19 |
| CD15 | 0.38 | 58.6 | 1.11 | CD61 | 0.014 | 13 | 4.92 |
| CD150 | 0.032 | 2.16 | 0.2 | CD62E | 0 | 0.00912 | 0.18 |
| CD151 | 26.9 | 99.6 | 98.5 | CD62L | 0.021 | 0.018 | 0.31 |
| CD152 | 0.011 | 0.029 | 0.47 | CD62P | 2.85 | 0.00891 | 0.27 |
| CD153 | 0.022 | 0.039 | 0.24 | CD63 | 99 | 99.8 | 96.7 |
| CD154 | 0.011 | 0.02 | 0.3 | CD64 | 0.014 | 0.00895 | 0.081 |
| CD158a | 0.032 | 0.021 | 0.25 | CD66 (a,c,d,e) | 63.4 | 23.3 | 35.7 |
| CD158b | 0 | 0.00983 | 0.16 | CD66b | 0.028 | 0 | 0.063 |
| CD15s | 0 | 0.018 | 0.18 | CD66f | 0.021 | 0.00911 | 0.21 |
| CD16 | 0.043 | 0 | 0.083 | CD69 | 0 | 0.0091 | 0.18 |
| CD161 | 0.032 | 0.019 | 0.21 | CD7 | 1.41 | 0.46 | 1.23 |
| CD162 | 0.021 | 0.039 | 0.34 | CD70 | 0.028 | 0.036 | 0.076 |
| CD163 | 0.032 | 0.02 | 0.4 | CD71 | 40.4 | 81.3 | 15.1 |
| CD164 | 66.1 | 77.9 | 70.8 | CD72 | 0.38 | 0.082 | 0.46 |
| CD165 | 99.8 | 99.9 | 91.9 | CD73 | 0.38 | 0.67 | 2.13 |
| CD166 | 100 | 30 | 94.9 | CD74 | 0.014 | 0.00934 | 0.12 |
| CD171 | 96.7 | 96.4 | 4.09 | CD75 | 0.16 | 0.064 | 18.3 |
| CD172b | 0.011 | 1.13 | 0.68 | CD77 | 0.21 | 0.11 | 0.45 |
| CD177 | 0 | 14.6 | 8.58 | cd79B | 0.021 | 0.00925 | 0.15 |
| CD178 | 7.9 | 0.02 | 0.39 | CD80 | 9.02 | 0.064 | 0.22 |
| CD18 | 0.011 | 0.041 | 0.17 | CD81 | 99.8 | 99.9 | 99.5 |
| CD180 | 0 | 0.00968 | 0.22 | CD83 | 0.029 | 0.037 | 1.1 |
| CD181 | 0 | 0.14 | 0.48 | CD84 | 0.028 | 0 | 0.21 |
| CD183 | 0.011 | 0.029 | 0.19 | CD85 | 0.016 | 0.00933 | 0.17 |
| CD184 | 0 | 99.9 | 5.62 | CD86 | 7.03 | 0.11 | 0.44 |
| CD19 | 0.021 | 0 | 0.12 | CD87 | 0.011 | 0 | 0.46 |
| CD193 | 0.085 | 0.27 | 0.52 | CD88 | 0.021 | 0.00944 | 0.25 |
| CD195 | 0.032 | 0.00998 | 0.37 | CD89 | 1.38 | 0 | 0.26 |
| CD196 | 0.053 | 0.059 | 0.36 | CD8a | 0.22 | 0.037 | 0.15 |
| CD197 | 7.41 | 0.03 | 0.34 | CD8b | 9.6 | 0.019 | 0.35 |
| CD1a | 0.021 | 0 | 0.34 | CD9 | 99.9 | 90.8 | 27.6 |
| CD1b | 0.021 | 0 | 0.2 | CD90 | 99.6 | 99.7 | 85.2 |
| CD1d | 0 | 0.05 | 0.16 | CD91 | 0.011 | 16 | 18.7 |
| CD2 | 0.032 | 0.01 | 0.15 | CD94 | 0.011 | 0 | 0.11 |
| CD20 | 0 | 0.036 | 0.18 | CD95 | 37.1 | 4.22 | 33 |
| CD200 | 100 | 99.9 | 99.3 | CD97 | 11.3 | 0.019 | 0.52 |
| CD201 | 0 | 0.39 | 53.1 | CD98 | 96.9 | 61.3 | 73.7 |
| CD205 | 55.3 | 0.17 | 4.85 | CD99 | 1.88 | 2.16 | 87 |
| CD206 | 0.011 | 0.04 | 0.23 | CD99R | 0.011 | 0.00952 | 1.37 |
| CD209 | 0.13 | 0.12 | 0.53 | CDw327 | 0.075 | 0.48 | 0.41 |
| CD21 | 58.2 | 0.095 | 0.38 | CDw328 | 0.011 | 0.00904 | 1.11 |
| CD210 | 0.011 | 0.019 | 0.26 | CDw329 | 0.16 | 0 | 0.59 |
| CD212 | 0 | 0.064 | 0.22 | CDw93 | 18 | 0.03 | 0.45 |
| CD22 | 0.032 | 0.00895 | 0.16 | CLIP | 0.011 | 0.036 | 0.3 |
| CD220 | 61 | 8.1 | 50.2 | CMRF-44 | 0.042 | 0.027 | 71.9 |
| CD221 | 99.9 | 100 | 92.6 | CMRF-56 | 0.011 | 0.00888 | 0.59 |
| CD226 | 0.074 | 0.02 | 0.34 | Cutaneous Lymph. Antigen | 0.011 | 0.046 | 0.78 |
| CD227 | 28 | 1.84 | 10.3 | Disialoganglioside GD2 | 0.18 | 9.34 | 1.21 |
| CD229 | 0.021 | 0.02 | 0.21 | EGF-r | 39.8 | 1.8 | 72.8 |
| CD23 | 23.1 | 0 | 0.31 | Fmlp-r | 0.085 | 0.027 | 1.79 |
| CD231 | 0 | 0 | 0.31 | gd TCR | 0.053 | 0.027 | 0.53 |
| CD235a | 0.011 | 0.18 | 0.41 | Hem. Prog. Cell | 27 | 1.79 | 20.5 |
| CD24 | 99.8 | 100 | 98 | HLA-A,B,C | 99.9 | 99.8 | 96.4 |
| CD243 | 0.021 | 0 | 0.55 | HLA-A2 | 91 | 20.2 | 26.4 |
| CD244 | 0.011 | 0.00994 | 0.73 | HLA-DQ | 14.4 | 8.09 | 10.9 |
| CD25 | 0.021 | 0 | 0.19 | HLA-DR | 2.41 | 0.22 | 0.69 |
| CD255 | 0.021 | 0.13 | 0.32 | HLA-DR,DP,DO | 0.095 | 0.00904 | 2.34 |
| CD26 | 0.19 | 1.71 | 37 | INT B7 | 0 | 0.00921 | 0.76 |
| CD267 | 0 | 0.046 | 0.27 | Invariant NKT | 0.053 | 0.00903 | 0.67 |
| CD268 | 0 | 0.02 | 0.41 | MIC A/B | 99.3 | 94.1 | 1.93 |
| CD27 | 0.011 | 0.0089 | 0.29 | mIgG1 | 0.032 | 0.045 | 0.52 |
| CD271 | 93 | 10.2 | 1.39 | mIgG2a | 0.053 | 0.018 | 0.66 |
| CD273 | 0.011 | 0.03 | 0.51 | mIgG2b | 0.053 | 0.036 | 0.38 |
| CD274 | 0 | 0.049 | 1.82 | mIgG3 | 0.064 | 0.00905 | 0.54 |
| CD275 | 0 | 1.06 | 1.03 | mIgM | 0.043 | 0.027 | 0.34 |
| CD278 | 0 | 0.02 | 0.51 | NKB1 | 0.011 | 0.0091 | 0.59 |
| CD279 | 5.89 | 0.098 | 1.11 | rlgG1 | 0 | 0.063 | 0.25 |
| CD28 | 0 | 0.018 | 0.13 | rlgG2a | 0.011 | 0.045 | 0.22 |
| CD282 | 0.075 | 0 | 0.55 | rlgG2b | 0.011 | 0.027 | 0.41 |
| CD29 | 99.5 | 93.4 | 70.7 | rlgM | 0 | 0.046 | 0.5 |
| CD294 | 0.011 | 0.036 | 0.32 | SSEA-1 | 17.8 | 86.9 | 13.6 |
| CD3 | 0.011 | 0.00958 | 0.2 | SSEA-3 | 0 | 7.7 | 0.55 |
| CD30 | 100 | 40.7 | 1.35 | SSEA-4 | 99.6 | 57.1 | 96.1 |
| CD305 | 0.13 | 0.027 | 0.38 | TRA-1-60 | 97.4 | 25.8 | 48.4 |
| CD309 | 91 | 42.9 | 1.41 | TRA-1-81 | 95.8 | 21 | 45.7 |
| CD31 | 0.27 | 0.046 | 1.34 | Vb 23 | 0.032 | 0.00891 | 0.4 |
|  |  |  |  | Vb 8 | 30.9 | 0.00889 | 0.29 |

**Table S2:**
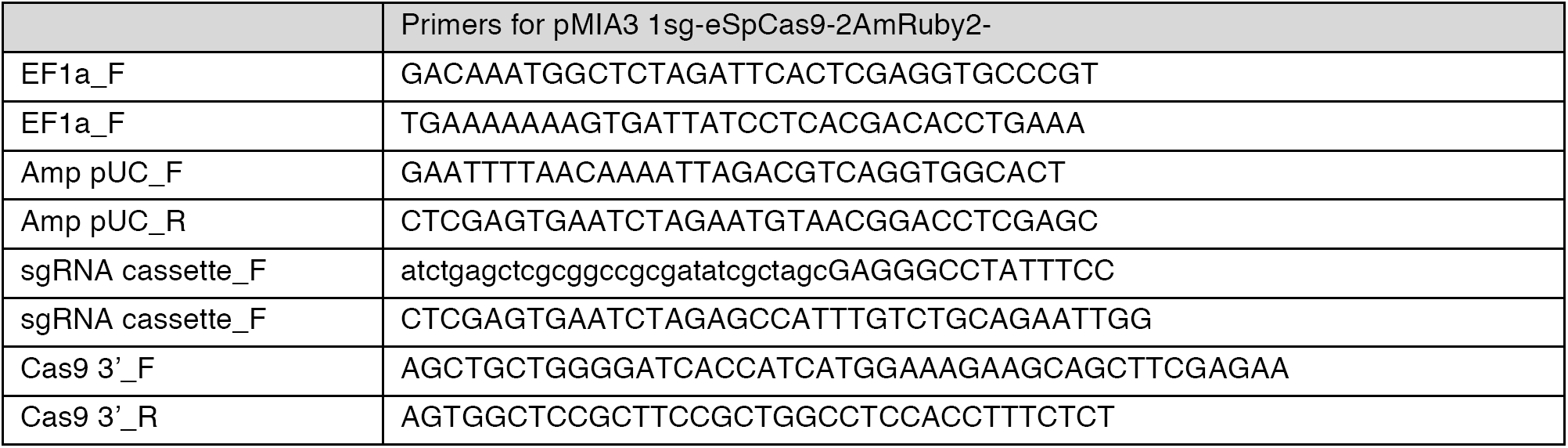
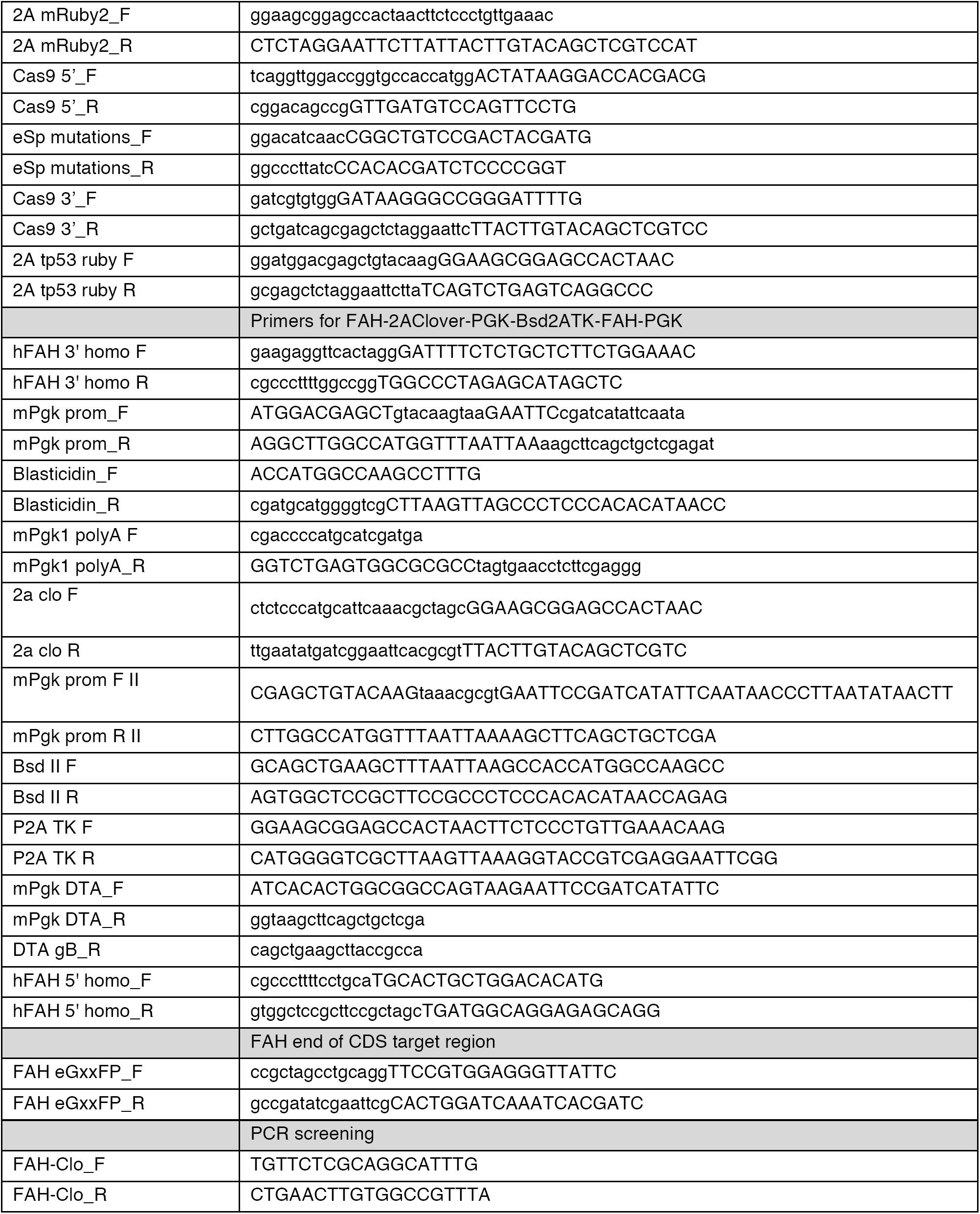
List of primers used for CRISPR-mediated gene editing

**Table S3:**
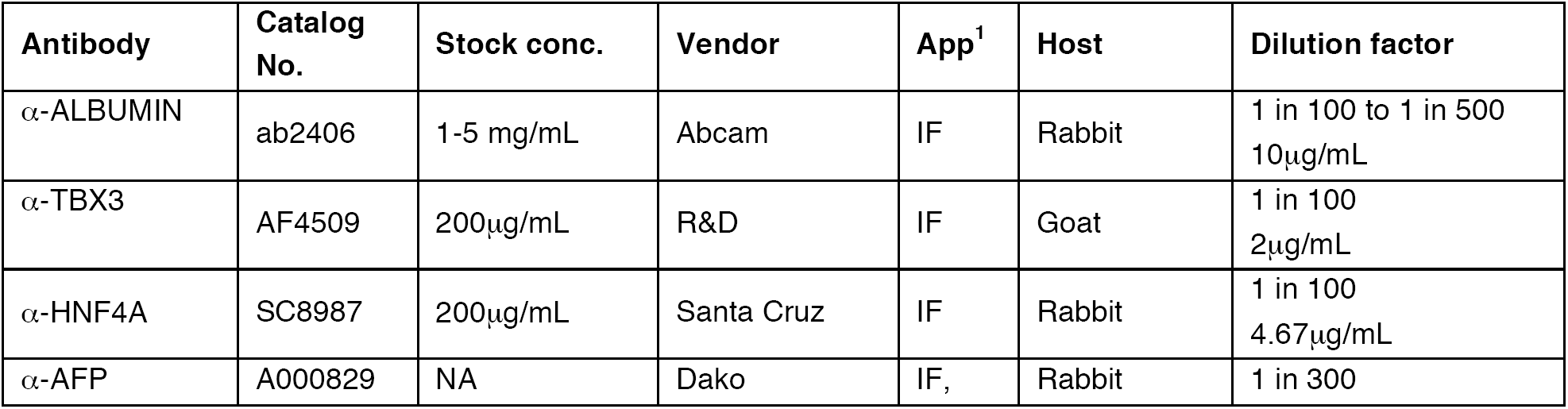
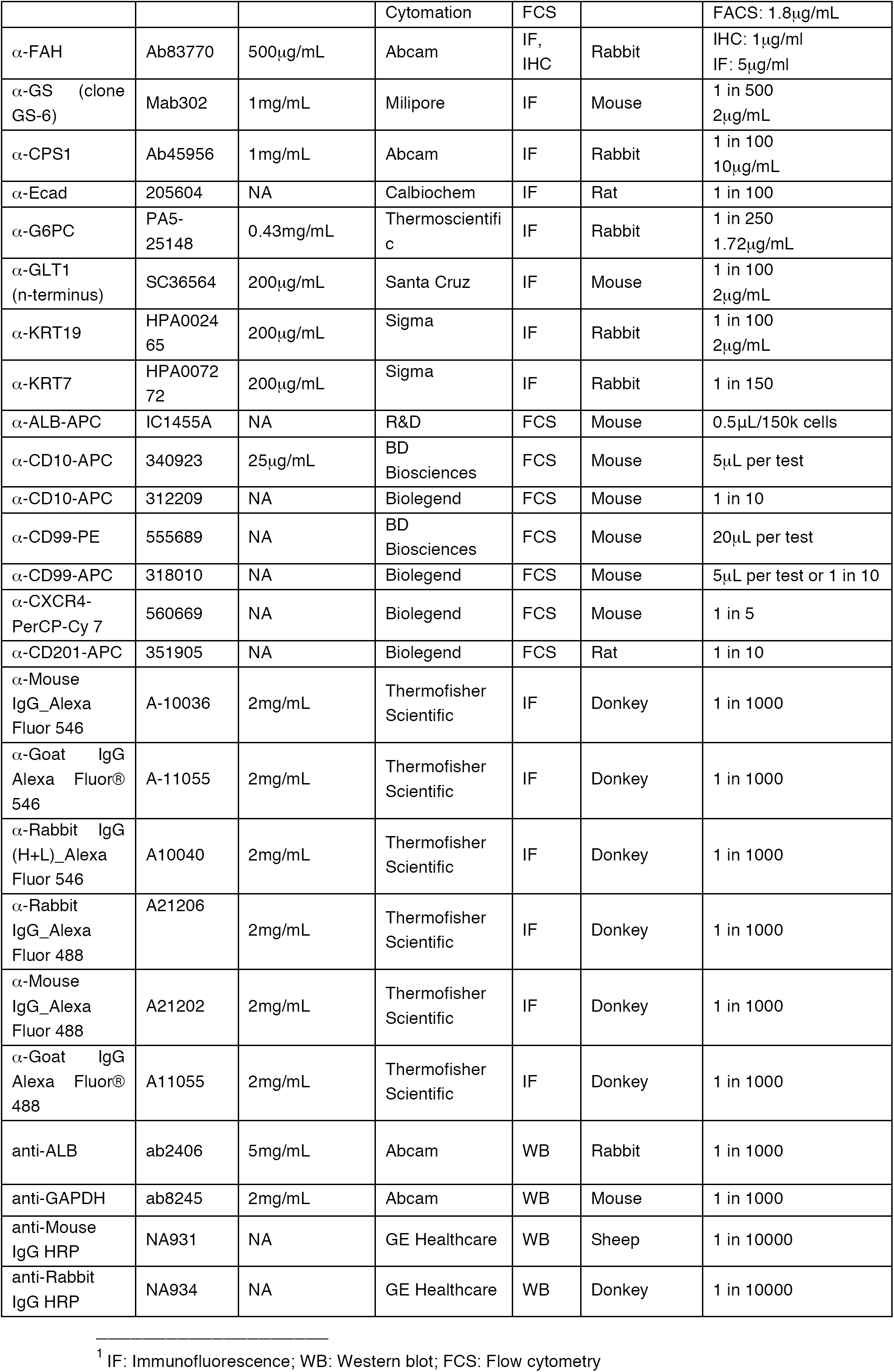
List of antibodies for immunostaining or FACS

| Antibody | Catalog No. | Stock conc. | Vendor | App<sup>1</sup> | Host | Dilution factor |
| --- | --- | --- | --- | --- | --- | --- |
| α-ALBUMIN | ab2406 | 1-5 mg/mL | Abcam | IF | Rabbit | 1 in 100 to 1 in 500<br>10μg/mL |
| α-TBX3 | AF4509 | 200μg/mL | R&D | IF | Goat | 1 in 100<br>2μg/mL |
| α-HNF4A | SC8987 | 200μg/mL | Santa Cruz | IF | Rabbit | 1 in 100<br>4.67μg/mL |
| α-AFP | A000829 | NA | Dako | IF, | Rabbit | 1 in 300 |
|  |  |  | Cytomation | FCS |  | FACS: 1.8µg/mL |
| α-FAH | Ab83770 | 500µg/mL | Abcam | IF,<br>IHC | Rabbit | IHC: 1µg/ml<br>IF: 5µg/ml |
| α-GS (clone<br>GS-6) | Mab302 | 1mg/mL | Milipore | IF | Mouse | 1 in 500<br>2µg/mL |
| α-CPS1 | Ab45956 | 1mg/mL | Abcam | IF | Rabbit | 1 in 100<br>10µg/mL |
| α-Ecad | 205604 | NA | Calbiochem | IF | Rat | 1 in 100 |
| α-G6PC | PA5-25148 | 0.43mg/mL | Thermoscientifi<br>c | IF | Rabbit | 1 in 250<br>1.72µg/mL |
| α-GLT1<br>(n-terminus) | SC36564 | 200µg/mL | Santa Cruz | IF | Mouse | 1 in 100<br>2µg/mL |
| α-KRT19 | HPA0024<br>65 | 200µg/mL | Sigma | IF | Rabbit | 1 in 100<br>2µg/mL |
| α-KRT7 | HPA0072<br>72 | 200µg/mL | Sigma | IF | Rabbit | 1 in 150 |
| α-ALB-APC | IC1455A | NA | R&D | FCS | Mouse | 0.5µL/150k cells |
| α-CD10-APC | 340923 | 25µg/mL | BD<br>Biosciences | FCS | Mouse | 5µL per test |
| α-CD10-APC | 312209 | NA | Biolegend | FCS | Mouse | 1 in 10 |
| α-CD99-PE | 555689 | NA | BD<br>Biosciences | FCS | Mouse | 20µL per test |
| α-CD99-APC | 318010 | NA | Biolegend | FCS | Mouse | 5µL per test or 1 in 10 |
| α-CXCR4-PerCP-Cy 7 | 560669 | NA | Biolegend | FCS | Mouse | 1 in 5 |
| α-CD201-APC | 351905 | NA | Biolegend | FCS | Rat | 1 in 10 |
| α-Mouse<br>IgG_Alexa<br>Fluor 546 | A-10036 | 2mg/mL | Thermofisher<br>Scientific | IF | Donkey | 1 in 1000 |
| α-Goat IgG<br>Alexa Fluor®<br>546 | A-11055 | 2mg/mL | Thermofisher<br>Scientific | IF | Donkey | 1 in 1000 |
| α-Rabbit IgG<br>(H+L)_Alexa<br>Fluor 546 | A10040 | 2mg/mL | Thermofisher<br>Scientific | IF | Donkey | 1 in 1000 |
| α-Rabbit<br>IgG_Alexa<br>Fluor 488 | A21206 | 2mg/mL | Thermofisher<br>Scientific | IF | Donkey | 1 in 1000 |
| α-Mouse<br>IgG_Alexa<br>Fluor 488 | A21202 | 2mg/mL | Thermofisher<br>Scientific | IF | Donkey | 1 in 1000 |
| α-Goat IgG<br>Alexa Fluor®<br>488 | A11055 | 2mg/mL | Thermofisher<br>Scientific | IF | Donkey | 1 in 1000 |
| anti-ALB | ab2406 | 5mg/mL | Abcam | WB | Rabbit | 1 in 1000 |
| anti-GAPDH | ab8245 | 2mg/mL | Abcam | WB | Mouse | 1 in 1000 |
| anti-Mouse<br>IgG HRP | NA931 | NA | GE Healthcare | WB | Sheep | 1 in 10000 |
| anti-Rabbit<br>IgG HRP | NA934 | NA | GE Healthcare | WB | Donkey | 1 in 10000 |

<sup>1</sup> IF: Immunofluorescence; WB: Western blot; FCS: Flow cytometry

**Table S4:**
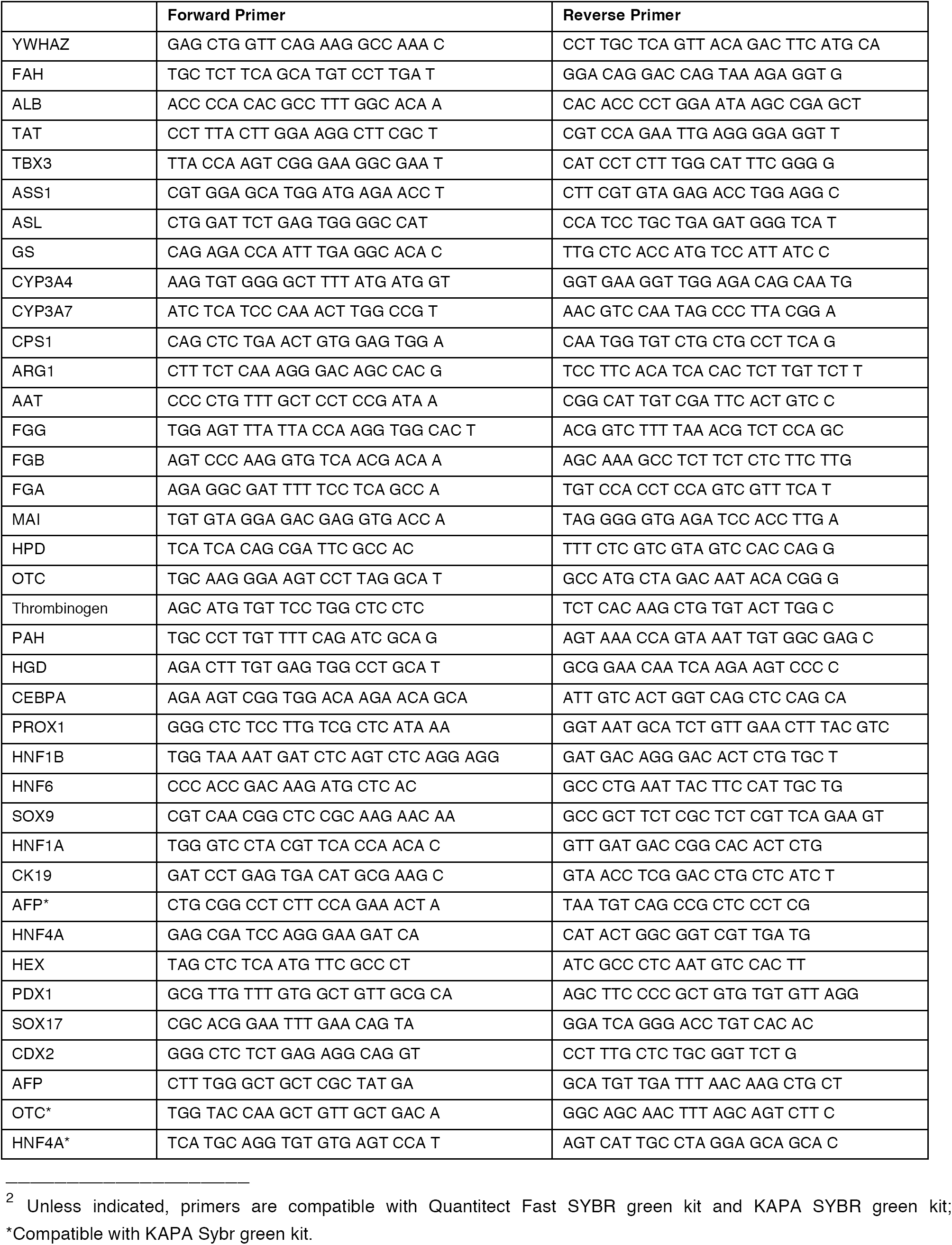
List of primers for qRT-PCR

## References

Agarwal, S., Holton, K.L., and Lanza, R. (2008). Efficient Differentiation of Functional Hepatocytes from Human Embryonic Stem Cells. Stem Cells 26, 1117–1127.

Azuma, H., Paulk, N., Ranade, A., Dorrell, C., Al-Dhalimy, M., Ellis, E., Strom, S., Kay, M.A., Finegold, M., and Grompe, M. (2007). Robust expansion of human hepatocytes in Fah-/-/Rag2-/-/Il2rg-/- mice. Nature Biotechnology 25, 903–910.

Basma, H., Gutiérrez, A.S., Yannam, G.R., Liu, L., Ito, R., Yamamoto, T., Ellis, E., Carson, S.D., Sato, S., Chen, Y., Muirhead, D., Álvarez, N.N., Wong, R.J., Chowdhury, J.R., Platt, J.L., Mercer, D.F., Miller, J.D., Strom, S.C., Kobayashi, N., and Fox, I.J. (2009). Differentiation and Transplantation of Human Embryonic Stem Cell–Derived Hepatocytes. YGAST 136, 990-999.e994.

Bayha, E., Jørgensen, M.C., Serup, P., and Grapin-Botton, A. (2009). Retinoic Acid Signaling Organizes Endodermal Organ Specification along the Entire Antero-Posterior Axis. PLoS One 4, e5845.

Bernardi, M., Maggioli, C., and Zaccherini, G. (2012). Human albumin in the management of complications of liver cirrhosis. Critical care (London, England) 16, 211.

Cai, J., Zhao, Y., Liu, Y., Ye, F., Song, Z., Qin, H., Meng, S., Chen, Y., Zhou, R., Song, X., Guo, Y., Ding, M., and Deng, H. (2007). Directed differentiation of human embryonic stem cells into functional hepatic cells. Hepatology 45, 1229–1239.

Carpentier, A., Tesfaye, A., Chu, V., Nimgaonkar, I., Zhang, F., Lee, S.B., Thorgeirsson, S.S., Feinstone, S.M., and Liang, T.J. (2014). Engrafted human stem cell–derived hepatocytes establish an infectious HCV murine model. Journal of Clinical Investigation 124, 4953–4964.

Cascio, S., and Zaret, K.S. (1991). Hepatocyte differentiation initiates during endodermal-mesenchymal interactions prior to liver formation. Development 113, 217–225.

Chambers, S.M., Qi, Y., Mica, Y., Lee, G., Zhang, X.-J., Niu, L., Bilsland, J., Cao, L., Stevens, E., Whiting, P., Shi, S.-H., and Studer, L. (2012). Combined small-molecule inhibition accelerates developmental timing and converts human pluripotent stem cells into nociceptors. Nature Biotechnology 30, 715–720.

Chung, W.-S., Shin, C.H., and Stainier, D.Y.R. (2008). Bmp2 Signaling Regulates the Hepatic versus Pancreatic Fate Decision. Developmental Cell 15, 738–748.

Cohen, D.E., and Melton, D. (2011). Turning straw into gold: directing cell fate for regenerative medicine. Nature Reviews Genetics 12, 243–252.

Colnot, S., and Perret, C. (2011). Liver Zonation. In (Boston, MA: Springer US), pp. 7–16.

D'amour, K.A., Agulnick, A.D., Eliazer, S., Kelly, O.G., Kroon, E., and Baetge, E.E. (2005). Efficient differentiation of human embryonic stem cells to definitive endoderm. Nature Biotechnology 23, 1534-1541.

Duncan, S.A. (2003). Mechanisms controlling early development of the liver. Mechanisms of Development 120, 19–33.

Espejel, S., Roll, G.R., McLaughlin, K.J., Lee, A.Y., Zhang, J.Y., Laird, D.J., Okita, K., Yamanaka, S., and Willenbring, H. (2010). Induced pluripotent stem cell–derived hepatocytes have the functional and proliferative capabilities needed for liver regeneration in mice. Journal of Clinical Investigation 120, 3120–3126.

Fukuda-Taira, S. (1981). Hepatic induction in the avian embryo: specificity of reactive endoderm and inductive mesoderm. Journal of embryology and experimental morphology 63, 111–125.

Gordillo, M., Evans, T., and Gouon-Evans, V. (2015). Orchestrating liver development. Development 142, 2094–2108.

Gouon-Evans, V., Boussemart, L., Gadue, P., Nierhoff, D., Koehler, C.I., Kubo, A., Shafritz, D.A., and Keller, G. (2006). BMP-4 is required for hepatic specification of mouse embryonic stem cell–derived definitive endoderm. Nature Biotechnology 24, 1402–1411.

Grapin-Botton, A. (2005). Antero-posterior patterning of the vertebrate digestive tract: 40 years after Nicole Le Douarin' PhD thesis. The International journal of developmental biology 49, 335–347.

Han, S. (2012). Generation of Functional Hepatic Cells from Pluripotent Stem Cells. Journal of Stem Cell Research and Therapy 01, 1–7.

Huang, P., Zhang, L., Gao, Y., He, Z., Yao, D., Wu, Z., Cen, J., Chen, X., Liu, C., Hu, Y., Lai, D., Hu, Z., Chen, L., Zhang, Y., Cheng, X., Ma, X., Pan, G., Wang, X., and Hui, L. (2014). Direct Reprogramming of Human Fibroblasts to Functional and Expandable Hepatocytes. Cell Stem Cell, 1-15.

Jung, J., Zheng, M., Goldfarb, M., and Zaret, K.S. (1999). Initiation of mammalian liver development from endoderm by fibroblast growth factors. Science 284, 1998–2003.

Jungermann, K. (1995). Zonation of metabolism and gene expression in liver. Histochemistry and Cell Biology 103, 81–91.

Karl, M.M., Howell, R.A., Hutchinson, J.H., and Catanzaro, F.J. (1953). Liver coma, with particular reference to management. AMA archives of internal medicine 91, 159–176.

Labelle, Y., Phaneuf, D., Leclerc, B., and Tanguay, R.M. (1993). Characterization of the human fumarylacetoacetate hydrolase gene and identification of a missense mutation abolishing enzymatic activity. Human Molecular Genetics 2, 941–946.

Lawson, K.A., Meneses, J.J., and Pedersen, R.A. (1991). Clonal analysis of epiblast fate during germ layer formation in the mouse embryo. Development 113, 891–911.

Ledouarin, N. (1964). Induction of prehepatic endoderm by mesoderm of the cardiac region in the chick embryo. Journal of embryology and experimental morphology 12, 651–664.

Lemaigre, F.P. (2009). Mechanisms of Liver Development: Concepts for Understanding Liver Disorders and Design of Novel Therapies. Gastroenterology 137, 62–79.

Lewis, S.L., and Tam, P.P.L. (2006). Definitive endoderm of the mouse embryo: formation, cell fates, and morphogenetic function. Developmental Dynamics 235, 2315–2329.

Loh, K.M., Ang, L.T., Zhang, J., Kumar, V., Ang, J., Auyeong, J.Q., Lee, K.L., Choo, S.H., Lim, C.Y.Y., Nichane, M., Tan, J., Noghabi, M.S., Azzola, L., Ng, E.S., Durruthy-Durruthy, J., Sebastiano, V., Poellinger, L., Elefanty, A.G., Stanley, E.G., Chen, Q., Prabhakar, S., Weissman, I.L., and Lim, B. (2014). Efficient Endoderm Induction from Human Pluripotent Stem Cells by Logically Directing Signals Controlling Lineage Bifurcations. Cell Stem Cell, 1–16.

Loh, K.M., Chen, A., Koh, P.W., Deng, T.Z., Sinha, R., Tsai, J.M., Barkal, A.A., Shen, K.Y., Jain, R., Morganti, R.M., Shyh-Chang, N., Fernhoff, N.B., George, B.M., Wernig, G., Salomon, R.E.A., Chen, Z., Vogel, H., Epstein, J.A., Kundaje, A., Talbot, W.S., Beachy, P.A., Ang, L.T., and Weissman, I.L. (2016). Mapping the Pairwise Choices Leading from Pluripotency to Human Bone, Heart, and Other Mesoderm Cell Types. Cell 166, 451–467.

McLin, V.A., Rankin, S.A., and Zorn, A.M. (2007). Repression of Wnt/-catenin signaling in the anterior endoderm is essential for liver and pancreas development. Development 134, 2207–2217.

Miyajima, A., Tanaka, M., and Itoh, T. (2014). Stem/Progenitor Cells in Liver Development, Homeostasis, Regeneration, and Reprogramming. Cell Stem Cell 14, 561–574.

Nissim, S., Sherwood, R.I., Wucherpfennig, J., Saunders, D., Harris, J.M., Esain, V., Carroll, K.J., Frechette, G.M., Kim, A.J., Hwang, K.L., Cutting, C.C., Elledge, S., North, T.E., and Goessling, W. (2014). Prostaglandin E2 Regulates Liver versus Pancreas Cell-Fate Decisions and Endodermal Outgrowth. Developmental Cell 28, 423–437.

Ogawa, S., Bear, C.E., Ahmadi, S., Chin, S., Li, B., Grompe, M., Keller, G., Kamath, B.M., Ogawa, M., and Ghanekar, A. (2015). Directed differentiation of cholangiocytes from human pluripotent stem cells. Nature Biotechnology 33, 853–861.

Ogawa, S., Surapisitchat, J., Virtanen, C., Ogawa, M., Niapour, M., Sugamori, K.S., Wang, S., Tamblyn, L., Guillemette, C., Hoffmann, E., Zhao, B., Strom, S., Laposa, R.R., Tyndale, R.F., Grant, D.M., and Keller, G. (2013). Three-dimensional culture and cAMP signaling promote the maturation of human pluripotent stem cell-derived hepatocytes. Development 140, 3285–3296.

Overturf, K., al-Dhalimy, M., Tanguay, R., Brantly, M., Ou, C.N., Finegold, M., and Grompe, M. (1996). Hepatocytes corrected by gene therapy are selected in vivo in a murine model of hereditary tyrosinaemia type I. Nature Genetics 12, 266–273.

Qi, Y., Zhang, X.-J., Renier, N., Wu, Z., Atkin, T., Sun, Z., Ozair, M.Z., Tchieu, J., Zimmer, B., Fattahi, F., Ganat, Y., Azevedo, R., Zeltner, N., Brivanlou, A.H., Karayiorgou, M., Gogos, J., Tomishima, M., Tessier-Lavigne, M., Shi, S.-H., and Studer, L. (2017). Combined small-molecule inhibition accelerates the derivation of functional cortical neurons from human pluripotent stem cells. Nature Biotechnology 35, 154–163.

Rashid, S.T., Corbineau, S., Hannan, N., Marciniak, S.J., Miranda, E., Alexander, G., Huang-Doran, I., Griffin, J., Ahrlund-Richter, L., Skepper, J., Semple, R., Weber, A., Lomas, D.A., and Vallier, L. (2010). Modeling inherited metabolic disorders of the liver using human induced pluripotent stem cells. Journal of Clinical Investigation 120, 3127–3136.

Rossi, J.M., Dunn, N.R., Hogan, B.L., and Zaret, K.S. (2001). Distinct mesodermal signals, including BMPs from the septum transversum mesenchyme, are required in combination for hepatogenesis from the endoderm. Genes & Development 15, 1998–2009.

Sampaziotis, F., de Brito, M.C., Madrigal, P., Bertero, A., Saeb-Parsy, K., Soares, F.A.C., Schrumpf, E., Melum, E., Karlsen, T.H., Bradley, J.A., Gelson, W.T.H., Davies, S., Baker, A., Kaser, A., Alexander, G.J., Hannan, N.R.F., and Vallier, L. (2015). Cholangiocytes derived from human induced pluripotent stem cells for disease modeling and drug validation. Nature Biotechnology 33, 845–852.

Sekiya, S., and Suzuki, A. (2011). Direct conversion of mouse fibroblasts to hepatocyte-like cells by defined factors. Nature 475, 390–393.

Sherwood, R.I., Maehr, R., Mazzoni, E.O., and Melton, D.A. (2011). Wnt signaling specifies and patterns intestinal endoderm. Mechanisms of Development 128, 387–400.

Shin, D., Shin, C.H., Tucker, J., Ober, E.A., Rentzsch, F., Poss, K.D., Hammerschmidt, M., Mullins, M.C., and Stainier, D.Y.R. (2007). Bmp and Fgf signaling are essential for liver specification in zebrafish. Development 134, 2041–2050.

Shiojiri, N., Wada, J.I., Tanaka, T., and Noguchi, M. (1995). Heterogeneous hepatocellular expression of glutamine synthetase in developing mouse liver and in testicular transplants of fetal liver. Laboratory Investigation 72, 740–747.

Shiraki, N., Umeda, K., Sakashita, N., Takeya, M., Kume, K., and Kume, S. (2008). Differentiation of mouse and human embryonic stem cells into hepatic lineages. Genes to Cells 13, 731–746.

Si Tayeb, K., Noto, F.K., Nagaoka, M., Li, J., Battle, M.A., Duris, C., North, P.E., Dalton, S., and Duncan, S.A. (2010). Highly efficient generation of human hepatocyte–like cells from induced pluripotent stem cells. Hepatology 51, 297–305.

Smith, D.D., and Campbell, J.W. (1988). Distribution of glutamine synthetase and carbamoyl-phosphate synthetase I in vertebrate liver. Proceedings of the National Academy of Sciences 85, 160–164.

Song, G., Pacher, M., Balakrishnan, A., Yuan, Q., Tsay, H.-C., Yang, D., Reetz, J., Brandes, S., Dai, Z., Pützer, B.M., Araúzo-Bravo, M.J., Steinemann, D., Luedde, T., Schwabe, R.F., Manns, M.P., Schöler, H.R., Schambach, A., Cantz, T., Ott, M., and Sharma, A.D. (2016). Direct Reprogramming of Hepatic Myofibroblasts into Hepatocytes In Vivo Attenuates Liver Fibrosis. Cell Stem Cell 18, 797–808.

Song, Z., Cai, J., Liu, Y., Zhao, D., Yong, J., Duo, S., Song, X., Guo, Y., Zhao, Y., Qin, H., Yin, X., Wu, C., Che, J., Lu, S., Ding, M., and Deng, H. (2009). Efficient generation of hepatocyte-like cells from human induced pluripotent stem cells. Cell Research 19, 1233–1242.

Spence, J.R., Lange, A.W., Lin, S.-C.J., Kaestner, K.H., Lowy, A.M., Kim, I., Whitsett, J.A., and Wells, J.M. (2009). Sox17 Regulates Organ Lineage Segregation of Ventral Foregut Progenitor Cells. Developmental Cell 17, 62–74.

Spence, J.R., Lauf, R., and Shroyer, N.F. (2011). Vertebrate intestinal endoderm development. Developmental Dynamics 240, 501–520.

Spence, J.R., Mayhew, C.N., Rankin, S.A., Kuhar, M.F., Vallance, J.E., Tolle, K., Hoskins, E.E., Kalinichenko, V.V., Wells, S.I., Zorn, A.M., Shroyer, N.F., and Wells, J.M. (2010). Directed differentiation of human pluripotent stem cells into intestinal tissue in vitro. Nature 470, 105–109.

Spijkers, J.A., van den Hoff, M.J., Hakvoort, T.B., Vermeulen, J.L., Tesink-Taekema, S., and Lamers, W.H. (2001). Foetal rise in hepatic enzymes follows decline in c-met and hepatocyte growth factor expression. Journal of Hepatology 34, 699–710.

St-Louis, M., and Tanguay, R.M. (1997). Mutations in the fumarylacetoacetate hydrolase gene causing hereditary tyrosinemia type I: overview. Human mutation 9, 291–299.

Stafford, D., and Prince, V.E. (2002). Retinoic acid signaling is required for a critical early step in zebrafish pancreatic development. Current biology : CB 12, 1215–1220.

Stanger, B.Z. (2015). Cellular Homeostasis and Repair in the Mammalian Liver. Annual review of physiology 77, 179–200.

Suzuki, A., Sekiya, S., Onishi, M., Oshima, N., Kiyonari, H., Nakauchi, H., and Taniguchi, H. (2008). Flow cytometric isolation and clonal identification of self-renewing bipotent hepatic progenitor cells in adult mouse liver. Hepatology 48, 1964–1978.

Takebe, T., Sekine, K., Enomura, M., Koike, H., Kimura, M., Ogaeri, T., Zhang, R.-R., Ueno, Y., Zheng, Y.-W., Koike, N., Aoyama, S., Adachi, Y., and Taniguchi, H. (2013). Vascularized and functional human liver from an iPSC-derived organ bud transplant. Nature, 1–5.

Tam, P.P., and Beddington, R.S. (1987). The formation of mesodermal tissues in the mouse embryo during gastrulation and early organogenesis. Development 99, 109–126.

Tennent, G.A., Brennan, S.O., Stangou, A.J., O'Grady, J., Hawkins, P.N., and Pepys, M.B. (2007). Human plasma fibrinogen is synthesized in the liver. Blood 109, 1971–1974.

Thomas, P.Q., Brown, A., and Beddington, R.S. (1998). Hex: a homeobox gene revealing peri-implantation asymmetry in the mouse embryo and an early transient marker of endothelial cell precursors. Development 125, 85–94.

Touboul, T., Hannan, N.R.F., Corbineau, S., Martinez, A., Martinet, C., Branchereau, S., Mainot, S., Strick-Marchand, H., Pedersen, R., Di Santo, J., Weber, A., and Vallier, L. (2010). Generation of functional hepatocytes from human embryonic stem cells under chemically defined conditions that recapitulate liver development. Hepatology 51, 1754–1765.

Tremblay, K.D., and Zaret, K.S. (2005). Distinct populations of endoderm cells converge to generate the embryonic liver bud and ventral foregut tissues. Developmental Biology 280, 87–99.

Wandzioch, E., and Zaret, K.S. (2009). Dynamic signaling network for the specification of embryonic pancreas and liver progenitors. Science 324, 1707–1710.

Yu, B., He, Z.-Y., You, P., Han, Q.-W., Xiang, D., Chen, F., Wang, M.-J., Liu, C.-C., Lin, X.-W., Borjigin, U., Zi, X.-Y., Li, J.-X., Zhu, H.-Y., Li, W.-L., Han, C.-S., Wangensteen, K.J., Shi, Y., Hui, L.-J., Wang, X., and Hu, Y.-P. (2013). Reprogramming Fibroblasts into Bipotential Hepatic Stem Cells by Defined Factors. Cell Stem Cell 13, 328–340.

Zhao, D., Chen, S., Duo, S., Xiang, C., Jia, J., Guo, M., Lai, W., Lu, S., and Deng, H. (2012). Promotion of the efficient metabolic maturation of human pluripotent stem cell-derived hepatocytes by correcting specification defects. Cell Research 23, 157–161.

Zhu, S., Rezvani, M., Harbell, J., Mattis, A.N., Wolfe, A.R., Benet, L.Z., Willenbring, H., and Ding, S. (2014). Mouse liver repopulation with hepatocytes generated from human fibroblasts. Nature, 1–18.

Zorn, A.M., and Wells, J.M. (2009). Vertebrate endoderm development and organ formation. Annual review of cell and developmental biology 25, 221–251.

## SUPPLEMENTAL REFERENCES

Abràmoff, M.D., and Magalhães, P.J. (2004). Image processing with Image. J Biophotonics International 11, 36–42.

Collins, T. (2007). ImageJ for microscopy. BioTechniques 43, S25–S30.

Cong, L., Ran, F.A., Cox, D., Lin, S., Barretto, R., Habib, N., Hsu, P.D., Wu, X., Jiang, W., Marraffini, L.A., and Zhang, F. (2013). Multiplex Genome Engineering Using CRISPR/Cas Systems. Science 339, 819–823.

Hockemeyer, D., Wang, H., Kiani, S., Lai, C.S., Gao, Q., Cassady, J.P., Cost, G.J., Zhang, L., Santiago, Y., Miller, J.C., Zeitler, B., Cherone, J.M., Meng, X., Hinkley, S.J., Rebar, E.J., Gregory, P.D., Urnov, F.D., and Jaenisch, R. (2011). Genetic engineering of human pluripotent cells using TALE nucleases. Nature Biotechnology 29, 731–734.

Huang, D.W., Sherman, B.T., and Lempicki, R.A. (2008). Systematic and integrative analysis of large gene lists using DAVID bioinformatics resources. Nature Protocols 4, 44–57.

Huang, D.W., Sherman, B.T., and Lempicki, R.A. (2009). Bioinformatics enrichment tools: paths toward the comprehensive functional analysis of large gene lists. Nucleic acids research 37, 1–13.

Ihry, R.J., Worringer, K.A., Salick, M.R., Frias, E., Ho, D., Theriault, K., Kommineni, S., Chen, J., Sondey, M., Ye, C., Randhawa, R., Kulkarni, T., Yang, Z., McAllister, G., Russ, C., Reece-Hoyes, J., Forrester, W., Hoffman, G.R., Dolmetsch, R., and Kaykas, A. (2017). P53 toxicity is a hurdle to CRISPR/CAS9 screening and engineering in human pluripotent stem cells. bioRxiv.

Kim, J.H., Lee, S.-R., Li, L.-H., Park, H.-J., Park, J.-H., Lee, K.Y., Kim, M.-K., Shin, B.A., and Choi, S.-Y. (2011). High Cleavage Efficiency of a 2A Peptide Derived from Porcine Teschovirus-1 in Human Cell Lines, Zebrafish and Mice. PLoS ONE 6, e18556.

Kleinstiver, B.P., Pattanayak, V., Prew, M.S., Tsai, S.Q., Nguyen, N.T., Zheng, Z., and Joung, J.K. (2016). High-fidelity CRISPR–Cas9 nucleases with no detectable genome-wide off-target effects. Nature 529, 490–495.

Lam, A.J., St-Pierre, F., Gong, Y., Marshall, J.D., Cranfill, P.J., Baird, M.A., McKeown, M.R., Wiedenmann, J., Davidson, M.W., Schnitzer, M.J., Tsien, R.Y., and Lin, M.Z. (2012). Improving FRET dynamic range with bright green and red fluorescent proteins. Nature Methods 9, 1005–1012.

Mashiko, D., Fujihara, Y., Satouh, Y., Miyata, H., Isotani, A., and Ikawa, M. (2013). Generation of mutant mice by pronuclear injection of circular plasmid expressing Cas9 and single guided RNA. Scientific Reports 3, 73.

Momcilovic, O., Choi, S., Varum, S., Bakkenist, C., Schatten, G., and Navara, C. (2009). Ionizing Radiation Induces Ataxia Telangiectasia Mutated-Dependent Checkpoint Signaling and G 2But Not G 1Cell Cycle Arrest in Pluripotent Human Embryonic Stem Cells. STEM CELLS 27, 1822–1835.

Momcilovic, O., Knobloch, L., Fornsaglio, J., Varum, S., Easley, C., and Schatten, G. (2010). DNA Damage Responses in Human Induced Pluripotent Stem Cells and Embryonic Stem Cells. PLoS ONE 5, e13410.

Reich, M., Liefeld, T., Gould, J., Lerner, J., Tamayo, P., and Mesirov, J.P. (2006). GenePattern 2.0. Nature Genetics 38, 500–501.

Slaymaker, I.M., Gao, L., Zetsche, B., Scott, D.A., Yan, W.X., and Zhang, F. (2015). Rationally engineered Cas9 nucleases with improved specificity. Science 351, 84–88.

Szymczak, A.L., Workman, C.J., Wang, Y., Vignali, K.M., Dilioglou, S., Vanin, E.F., and Vignali, D.A.A. (2004). Correction of multi-gene deficiency in vivo using a single ' self-cleaving' 2A peptide– based retroviral vector. Nature Biotechnology 22, 589–594.

